# Ecdysone exerts biphasic control of regenerative signaling, coordinating the completion of regeneration with developmental progression

**DOI:** 10.1101/2021.08.12.456119

**Authors:** Faith Karanja, Subhshri Sahu, Sara Weintraub, Rajan Bhandari, Rebecca Jaszczak, Jason Sitt, Adrian Halme

## Abstract

In *Drosophila melanogaster*, loss of regenerative capacity in wing imaginal discs coincides with an increase in systemic levels of the steroid hormone ecdysone, a key coordinator of their developmental progression. Regenerating discs release the relaxin hormone Dilp8, which limits ecdysone synthesis and extends the regenerative period. Here, we describe how regenerating tissues produce a biphasic response to ecdysone levels: lower concentrations of ecdysone promote local and systemic regenerative signaling, whereas higher concentrations suppress regeneration through the expression of *broad* splice isoforms. Ecdysone also promotes the expression of *wingless* during both regeneration and normal development through a distinct regulatory pathway. This dual role for ecdysone explains how regeneration can still be completed successfully in *dilp8*^-^ mutant larvae: higher ecdysone levels increase the regenerative activity of tissues, allowing regeneration to reach completion in a shorter time. From these observations, we propose that ecdysone hormone signaling functions to coordinate regeneration with developmental progression.

**Summary Statement:** Ecdysone coordinates regenerative activity with developmental progression through the biphasic, concentration-dependent activation, and suppression of regenerative signaling.

## Introduction

As tissues develop, their capacity to regenerate is often diminished (Seifert & Voss, 2013; Yun, 2015). In many cases, loss of regenerative capacity is developmentally regulated and coincides with changes in systemic hormone signaling. For example, loss of regenerative capacity in the heart tissues of both *Xenopus laevis* and mice is preceded by a sharp increase in systemic thyroid hormone levels (Hirose et al., 2019; Marshall et al., 2019). Similarly, *Drosophila melanogaster* imaginal discs (the larval precursors to adult tissues) lose the ability to regenerate near the end of larval development (Halme et al., 2010), coinciding with an increase in systemic levels of the steroid hormone ecdysone, a key coordinator of *Drosophila* developmental progression (Burdette, 1962; Yamanaka et al., 2013). Although thyroid hormone in vertebrates and ecdysone in *Drosophila* have also been associated with loss of regenerative capacity, both hormones are present at lower levels during the regeneration-competent periods of development (Hirose et al., 2019; Hodgetts et al., 1977; Lavrynenko et al., 2015; Marshall et al., 2019). The regulation of systemic levels of ecdysone is a crucial part of the *Drosophila* regenerative response. Regenerating imaginal discs synthesize and release the relaxin hormone *Drosophila* insulin-like peptide 8 (Dilp8), which signals to the brain and endocrine organs through its receptor Lgr3 to limit the synthesis of the steroid hormone ecdysone (Colombani et al., 2012, 2015; Garelli et al., 2012, 2015; Jaszczak et al., 2016; Vallejo et al., 2015). Reduced ecdysone production extends the larval developmental period, providing damaged imaginal discs additional time to regenerate (Halme et al., 2010). However, it is unclear whether low levels of ecdysone still can influence regenerative activity.

To better understand how ecdysone regulates regeneration, we examined how manipulating ecdysone signaling affects both systemic and local regenerative pathways in the *Drosophila* wing imaginal disc. Here we demonstrate that while ecdysone signaling is limited during regeneration, it remains necessary for the regenerative response. We show that ecdysone signaling limits regeneration at the end of larval development through the expression of specific splice isoforms of the BTB-POZ transcription factor Broad, which inhibit the regenerative expression of Wingless (Wg). However, we also establish that ecdysone signaling is essential for regenerative activity and Wg expression in the disc. Regenerating discs exhibit a positive response in signaling activity to increasing ecdysone levels promoting Wg expression through a Broad-independent pathway. This dual role for ecdysone in promoting and limiting regeneration helps explain how *dilp8^-^* mutant larvae, which lack the regenerative checkpoint and thus produce minimal developmental delay following damage, can still regenerate their wing discs during the shorter regenerative period. Therefore, ecdysone’s biphasic regulation of regenerative activity gives *Drosophila* larvae the ability to coordinate the completion of regeneration with the end of the larval period.

## Results

### Ecdysone limits regenerative repair of wing imaginal discs

To examine how ecdysone signaling regulates regenerative activity, we measured how developmental timing and changes in ecdysone titer regulate regenerative outcomes following X-irradiation damage to wing imaginal discs. *Drosophila* larvae exposed to X-irradiation during early 3^rd^ larval instar (80h AED @ 25°C) can regenerate their wing tissues almost entirely, with only a few adult wings from irradiated larvae exhibiting minor defects (Fig. 1A, B, Fig S1A). The regenerated adult wings are also a similar size to those produced by undamaged control larvae (Fig. 1C). In contrast, larvae irradiated at a later, pre-pupal stage (late 3^rd^ larval instar – 104h AED @ 25°C) produced adult wings that exhibit a greater frequency of malformations in the wing veins and margin (Fig. 1A, B) and fail to reach the size of normal undamaged wings (Fig. 1C). These differences in regenerative capacity are also correlated with the ability to activate the regenerative checkpoint. Irradiation of larvae at 80h AED produces a robust checkpoint activation and developmental delay, whereas irradiation at 104h AED fails to activate the regeneration checkpoint, producing no extension of the larval period (Fig 1D). These results are consistent with previous observations that identified this developmental transition as a regeneration restriction point (RRP), a developmental period when damage no longer activates the regenerative checkpoint and tissues lose their regenerative capacity (Halme et al., 2010; Smith-Bolton et al., 2009). To examine how transition through the RRP impacts regenerative signaling in damaged wing discs, we measured the damage-induced expression of *dilp8*. Dilp8 is a critical regulator of the systemic response to regeneration, an effector of the regeneration checkpoint, and a valuable marker for regenerative activity in damaged tissues. Consistent with the reduced regenerative activity we observed as larvae pass through the RRP, we observe reduced activation of *dilp8* expression (Dilp8::GFP) in wing discs damaged at progressively later times in larval development (Fig. 1E and S1B, C).

**Figure 1.**
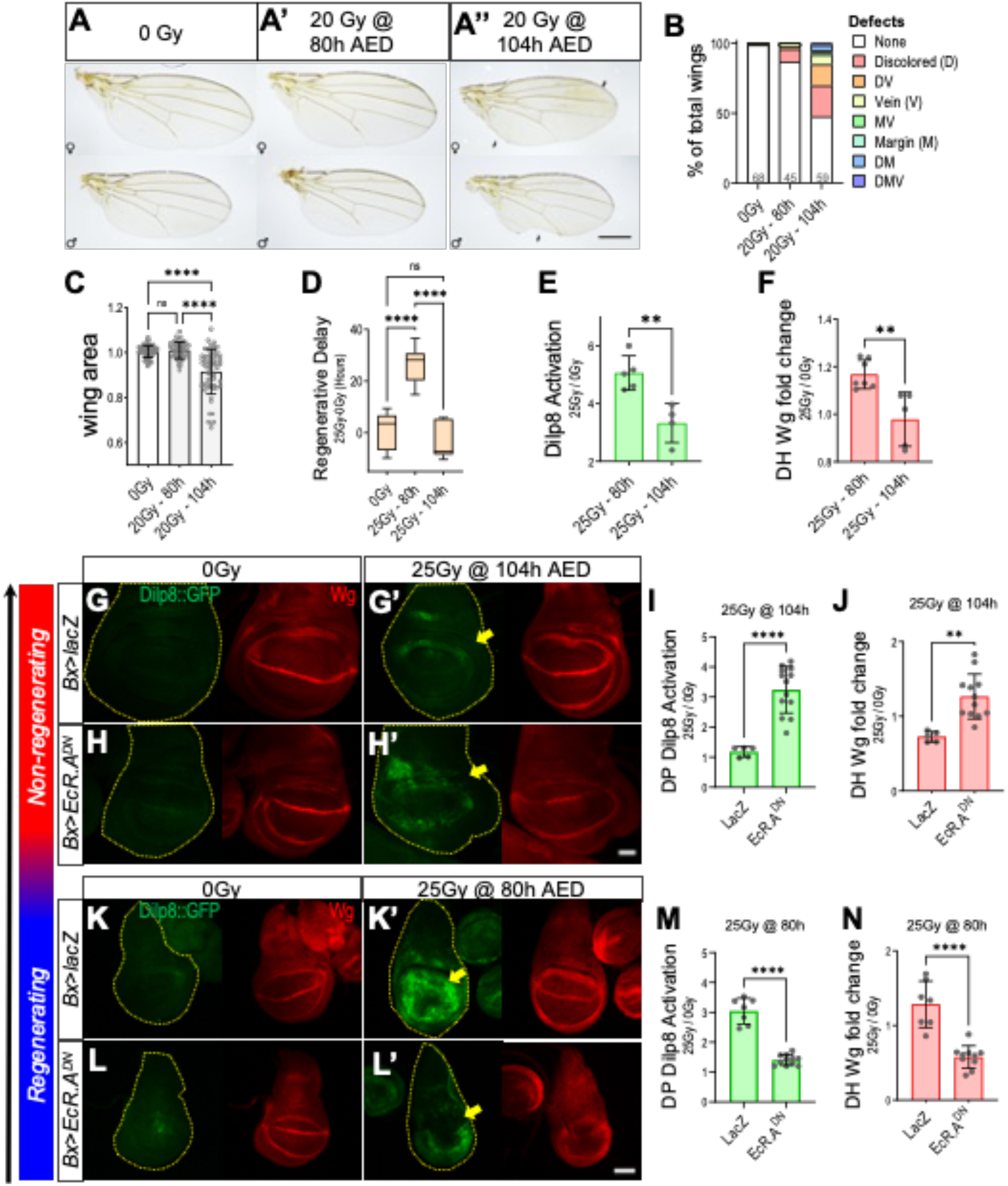
Ecdysone signaling is necessary for the suppression and activation of regeneration pathways. **(A)** Representative images of adult female and male wings isolated from flies that were undamaged (A), damaged in early 3^rd^ larval instar - 20Gy X-irradiation at 80h AED (A’), or damaged in late 3^rd^ larval instar - 20Gy X-irradiation at 104h AED, when regenerative capacity is restricted (A’’). Black arrows indicate defects on late damage wings. Scale = 500cm. **(B)** Adult wings that show individual defects or combinations of defects increase following late damage. The graph shows the percentage of defective adult wings from larvae that had no damage (0Gy), early damage (25Gy-80h), and late damage (20Gy-104h) during wing development. Population size is indicated in the graph. **(C)** Quantification of adult wing size following no damage (0Gy), early damage (20Gy-80h), and late damage (20Gy-104h). Size of wing measured in the unit area and normalized to undamaged wing size of respective sex. One-way ANOVA with Tukey’s multiple comparisons test, ****p<0.0001. **(D)** Quantification of regenerative delay in undamaged (0Gy), early damage (20Gy-80h), and late damage (20Gy-104h). One-way ANOVA with Tukey’s multiple comparisons test, ****p<0.0001. **(E-F)** Quantification of relative Dilp8::GFP expression in wing pouch (E) and Wg expression in Dorsal Hinge (DH) (F); normalized to expression levels in undamaged tissues at each respective timepoint tissues. **p<0.01, Unpaired t-test. **(G-H)** Representative images of pouch Dilp8::GFP (green) and dorsal hinge Wg (red) expression in 116h AED wing imaginal discs. The yellow dotted line indicates tissue area. Tissues are expressing lacZ (*bx>lacZ*) as a control (G-G’) or EcR.A^DN^ (*bx>EcR.A^W650A^*, H-H’) in the dorsal wing pouch region, indicated by yellow arrows. Tissues were either left undamaged (G and H) or damaged late, 25Gy@104h AED (G’ and H’), then isolated 12 hours after damage timepoint. Scale bar = 50um. **(I-J)** Quantification of relative regenerative activity using fold change in Dilp8::GFP expression in the dorsal wing pouch (DP) (I) and dorsal hinge (DH) Wg expression (J) following late damage (25Gy-104h), in *bx>lacZ* and *bx>EcR.A^DN^* wing imaginal discs. Fold change determined by normalizing to respective undamaged tissues, ****p<0.0001, **p<0.01, Mann Whitney t-test. **(K-L)** Representative images of pouch Dilp8::GFP (green) and dorsal hinge Wg (red) expression in 92h AED wing imaginal discs. The yellow dotted line indicates tissue area. Tissues are expressing lacZ (K-K’) or *EcR.A^DN^* (L-L’) in the dorsal wing pouch region, indicated by yellow arrows. Tissues were either left undamaged (K and L) or damaged early, 25Gy@80h AED (K’ and L’), then isolated 12 hours after damage timepoint. Scale bar = 50um. **(M-N)** Quantification of relative regenerative activity using fold change in Dilp8::GFP expression in the dorsal wing pouch (M) and dorsal hinge (DH) Wg expression (N) following early damage (25Gy-80h), in *Bx>lacZ* and *Bx>EcR.A^DN^* wing imaginal discs. Fold change determined by normalizing to respective undamaged tissues, ****p<0.0001, **p<0.01, *p<0.05, Mann Whitney t-test.

To further examine whether regenerative signaling in damaged discs is limited as larvae pass through the RRP, we also examined the irradiation-induced expression of the critical regenerative morphogen, Wingless (Wg). Wg is upregulated in the regeneration blastema in damaged wing discs and is necessary for regeneration (Smith-Bolton et al., 2009). Wg expression in the hinge region surrounding the wing pouch is critical to the radiation-resistant cells in this region that contribute to the X-irradiation regenerative response (Verghese & Su, 2016). When we examine Wg expression in the dorsal hinge of irradiated larvae, we find that early irradiation (80h AED) produces a significant increase in Wg expression, which is no longer observed as larvae transit the RRP (Fig. 1F, S1D, E). These results demonstrate that the loss of regenerative activity seen as larvae transit the RRP is accompanied by a failure to activate the regenerative checkpoint and the inability to activate the expression of Dilp8 and Wg, key mediators of systemic and local regenerative processes.

As larvae approach the larval/pupal transition, circulating levels of ecdysone increase rapidly and promote the exit from the larval period (Lavrynenko et al., 2015; Rewitz et al., 2013). To determine whether increased ecdysone titer is sufficient to limit regenerative activity in wing discs, ecdysone levels were increased in larvae ectopically by feeding larva food containing 20-hydroxyecdysone (20HE), an active form of this steroid hormone. Larvae damaged before the RRP (80h AED) and fed 0.3 mg/ml 20HE no longer completely regenerate their imaginal discs (Fig. S2A,B) but instead produce malformed (Fig. S2C) and smaller (Fig.S2D) adult wings. Furthermore, feeding low levels of 20HE (0.1mg/ml) to larvae irradiated after the RRP (104h AED) produces a synergistic increase in adult wing malformations (Fig. S2E,F,G) and suppression of regenerative growth (Fig. S2H). Together these observations support a model that the increasing levels of systemic ecdysone signaling at the end of larval development suppress regenerative signaling and growth in wing imaginal discs.

### Ecdysone signaling in the wing disc is necessary for both the suppression and activation of regenerative signals

Although the 20HE feeding experiments above demonstrate that increasing systemic ecdysone limits the regeneration observed in adult wing tissues, it remained unclear whether ecdysone signaling acts directly on regenerating tissues to suppress regenerative activity or indirectly through other tissues. To test the tissue-autonomous requirement for ecdysone signaling in regenerating wing discs, we expressed a dominant-negative allele of the ecdysone receptor (Cherbas et al., 2003) in the dorsal compartment of the wing pouch using *Beadex*-driven, Gal4-UAS expression (*Bx>EcR.A^DN^*, Fig. S3A). After larvae transition through the RRP (104h AED), regeneration-induced expression of the Dilp8 checkpoint signal is limited (Fig. 1G,I, S3B), reflecting the reduced regenerative activity in these tissues. However, we see that targeted inhibition of ecdysone signaling in the dorsal wing pouch significantly increases *dilp8* expression in larvae damaged at 104h AED (Fig. 1H,I, S3B), suggesting that the regenerative Dilp8 expression in these post-RRP tissues is limited by ecdysone signaling.

To further examine whether regenerative signaling is increased in damaged post-RRP wing discs when we limit ecdysone signaling, we also examined the damage-induced expression of Wg at the dorsal hinge region of the wing pouch (Fig. S3A). Prior to the RRP, when discs are competent to regenerate, we observe that damage induces an increase in Wg expression at the dorsal hinge (Fig. 1F, S1D,E). Inhibition of ecdysone signaling in the dorsal pouch leads to an overall decrease in Wg expression at the hinge in undamaged tissues (Fig. 1G, S3C), an observation we address more specifically later in this study. However, in contrast to control discs, where no significant increase in Wg expression is seen in the dorsal hinge of discs damaged after the RRP (104h AED), we see that limiting ecdysone signaling in the wing discs now permits a damage-induced increase in dorsal hinge Wg expression in post-RRP wing discs. This increase in Wg expression is similar to what we see in regeneration competent discs pre-RRP (Fig. 1H, J, S3C). These data demonstrate that at the end of larval development, ecdysone signaling acts tissue-autonomously in wing discs to suppress critical local (Wg upregulation at the hinge) and systemic (*dilp8* expression) signaling events associated with regeneration.

Since circulating ecdysone is present at lower levels at earlier stages of larval development when the wing discs can regenerate (Lavrynenko et al., 2015), we wanted to determine whether ecdysone regulates regeneration signaling in wing discs damaged at 80h AED, before the RRP. To assess this, we X-irradiated control and *Bx>EcR.A^DN^* larvae early in the third larval instar (80h AED), when regenerative activity is high. Unexpectedly, we observe that ecdysone signaling is necessary for the activation of regenerative signaling pathways following early damage. There is a clear inhibition of *dilp8* expression in the regenerating dorsal wing of *Bx>EcR.A^DN^* larvae compared to the controls (Fig. 1K, L, M, S3D). Ecdysone signaling is also necessary for the increased expression of Wg in the dorsal hinge following damage as we observed reduced expression of Wg at the dorsal hinge of *Bx>EcR.A^DN^* expressing tissue compared with controls [Fig. 1K,L,N, S3E]. This requirement of ecdysone signaling in the activation of regenerative activity is also seen following targeted expression of the *Drosophila* TNFa homolog, *eiger* (*Bx>egr*). Overexpression of *eiger* in wing discs produces localized damage and elicits a strong induction of Wg and Dilp8 expression in the regeneration blastema ((Smith-Bolton et al., 2009), Fig. S3F). We see that expression of *EcR.A^DN^* decreases *eiger*-induced Wg and Dilp8 expression in the damage blastema formed in the dorsal compartment of the wing pouch (Fig. S3F-H).

Together, these findings suggest a dual (activation and suppression) role for ecdysone signaling in the regulation of regenerative activity. During regenerative competence, ecdysone signaling in the damaged disc is required to activate Wg and Dilp8 expression, two critical signaling events that coordinate the local and systemic regenerative responses, respectively. Following development past the RRP, when the imaginal discs lose their regenerative capacity, ecdysone signaling in the disc is required to suppress the activation of these regenerative pathways.

### Ecdysone regulates regenerative signaling in a biphasic, concentration-dependent manner

During the last larval instar, pulses of ecdysone synthesis increase the systemic levels of circulating ecdysteroids in the larvae before a final surge of ecdysone synthesis at the end of larval development activates pupariation pathways and initiate metamorphosis (Lavrynenko et al., 2015). Based on this, we hypothesized that the dual activities of ecdysone signaling that we had observed, being necessary for activation of regenerative pathways during regenerative competence and suppressing regenerative pathways following development past the RRP, could reflect the different ecdysone levels at these two points of development. Lower circulating concentrations of ecdysone, such as those found pre-RRP, are necessary for activation of regenerative activity, whereas the higher levels of ecdysone circulating during the pre-pupal surge interfere with the activation of regeneration pathways. To test this hypothesis, we manipulated circulating 20HE levels in larvae by supplementing their food with increasing concentrations of 20HE following X-irradiation damage at 80h AED. We then measured the regenerative activation Wg and Dilp8 expression in wing discs 12 hours after X-irradiation (Fig. 2A).

**Figure 2.**
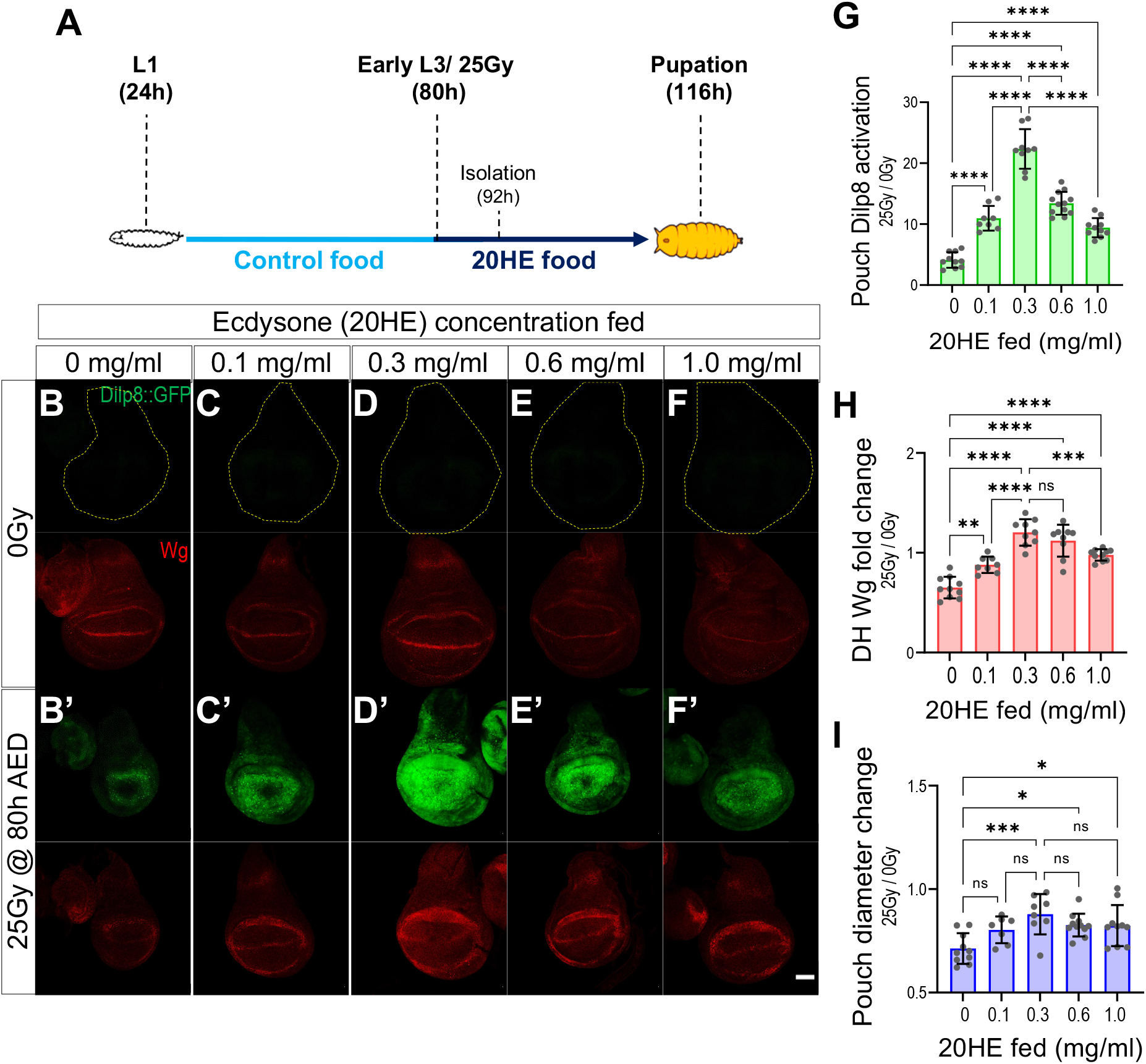
Ecdysone regulates regenerative signaling in a biphasic, concentration-dependent manner. **(A)** Schematic of ecdysone (20HE in ethanol) feeding experiment. *w^1118^* larvae were fed various 20HE concentrations in early third instar (80h AED) immediately after irradiation damage (25Gy @ 80h AED) or no damage (0Gy). Tissues were isolated 12 hours after damage and feeding (92h AED). **(B-F)** Representative images of pouch Dilp8::GFP (green) and dorsal hinge Wg (red) expression in 92h AED undamaged (B-F) and early damaged – 25Gy @ 80h AED (B’-F’) wing imaginal discs. The yellow dotted line indicates tissue area. The *w^1118^* larvae were fed 20HE of various concentrations, 0mg/ml (B-B’), 0.1mg/ml (C-C’), 0.3mg/ml (D-D’), 0.6mg/ml (E-E’) and 1.0mg/ml (F-F’). Scale bar = 50um. **(G-I)** Quantification of relative regenerative activity using fold change of Dilp8::GFP expression in the wing pouch (G), dorsal hinge (DH), Wg expression (H), and pouch diameter (I) in 20HE fed and early damaged *w^1118^* wing imaginal discs. Fold change determined by normalizing to respective undamaged tissues, ****p<0.0001, ***p<0.001, **p<0.01, *p<0.05, One-way ANOVA with Tukey’s multiple comparisons tests

We find that feeding larvae ecdysone generally promotes activation of regeneration genes following X-irradiation damage. At all concentrations of 20HE feeding, we observe an increase in Dilp8 and Wg expression in damaged wing discs compared with control larvae with no 20HE supplement in their food (Fig. 2B-F, G, H S4A, B). We also observe that 20HE feeding increased the size of the regenerating tissue (Fig. 2I, S4C). However, the effect of ecdysone feeding was maximized at 0.3 mg/ml 20HE concentration. Higher concentrations of ecdysone (0.6 or 1.0 mg/ml) produce a substantial reduction in Dilp8 and Wg expression (Fig. 2B-F, G, H S4A, B) and produce no additional increase in growth (Fig. 2I, S4C). Therefore, 20HE feeding produces a biphasic regenerative signaling response in irradiated tissues. In contrast, the biphasic effect of increasing 20HE concentrations through feeding is not seen when we examine *eiger*-induced Dilp8 and Wg expression, with both showing a modest but not statistically significant increase in expression with increasing ecdysone levels (Fig S4D-F). The differences in ecdysone sensitivity from X-irradiated tissues may reflect the persistence and the intensity of the damage produced in the *Bx>eiger* tissues, which likely maximizes regenerative signaling.

### Broad splice isoforms are necessary to block regenerative signaling after the RRP

To determine how ecdysone limits regenerative signaling, we examined the expression of one of the downstream targets of ecdysone signaling, the BTB-POZ family transcription factor, Broad. Broad is one of the earliest targets of the pre-pupal ecdysone pulse. Splice isoforms of the transcription factor *broad* – (*brZ1, Z2, Z3, and Z4*), named after their respective zinc finger domains ((Bayer et al., 1996; DiBello et al., 1991; Kiss et al., 1988); Fig. S5A), determine the tissue-specific signaling events that are initiated in response to ecdysone (Emery et al., 1994; Von Kalm et al., 1994). BrZ1 has also been recently shown to antagonize Chinmo expression in the wing disc, limiting regenerative activity (Narbonne-Reveau & Maurange, 2019). Using western-blotting of wing disc-derived lysates (WB, Fig. 3A) and immunofluorescence with Broad-targeting antibodies (IF, Fig. S5B), we visualized the spatial and temporal distribution of Broad expression during normal wing development. Broad splice isoforms are expressed in all imaginal disc cells throughout the final instar of larval development. Based on their distinct molecular sizes (Emery et al., 1994), we determined that BrZ2 is expressed throughout the third larval instar, but its levels increase as the larvae approach pupation. While we could not distinguish BrZ1 and BrZ3 based on size, previous studies have demonstrated that the BrZ3 splice isoform is not expressed in imaginal discs (Emery et al., 1994). The expression of both BrZ1 and BrZ4 can be detected at 104hAED and dramatically increase as the larvae approach pupariation. We could verify the emergence of BrZ1 expression in the tissue using Z1-targeted IF (Fig. S5B’). Following early damage (2.5kR@80h AED), the expression of the Broad isoforms is delayed (Fig. 3A). Therefore, the expression of Broad isoforms corresponds to the known changes in ecdysone levels during larval development and regeneration, and increased Broad expression correlates with the loss of regenerative capacity.

**Figure 3.**
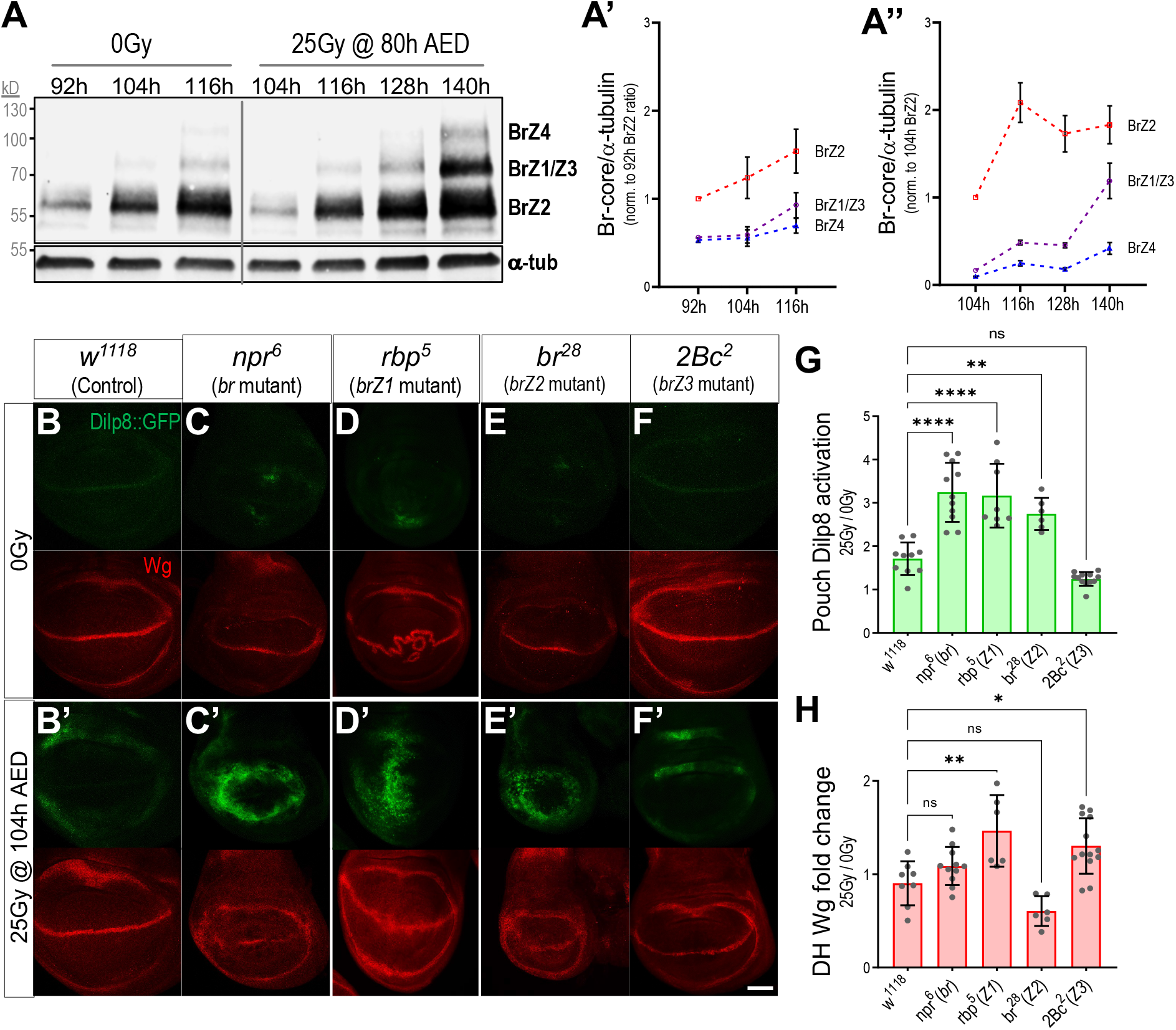
Broad isoforms are necessary for the restriction of regenerative activity at the end of larval development. **(A)** Western Blot (A) time course of Broad isoform expression in undamaged and early damaged *w^1118^* wing imaginal discs. Tissues were isolated in 12-hour intervals. Due to limitations of tissue size, isolation of tissues started at 92h AED for undamaged tissues and 104h AED for damaged tissues. Broad core antibody was used to visualize BrZ4 (~110kD), BrZ1/3 (~90kD), and BrZ2 (~55-65kD), while and a-tub (~50kD) was used as a loading control. Protein size (kD) ladder - on the left. Quantification of Broad expression in undamaged (A’) normalized to 92h AED BrZ2 expression, n=5. Quantification of broad expression in damaged (A’’) wing discs normalized to 104h AED BrZ2 expression, n=3. **(B-F)** Loss of Broad isoforms allows for activation of regenerative activity past the regeneration restrictive time point. Images show representative examples of pouch Dilp8::GFP (green) and dorsal hinge Wg (red) expression in 116h AED wing discs of undamaged (B-F), and late damage – 25Gy @ 104h AED (B’-F’), control – *w^1118^* (B-B’), and *br* mutants: full *br* mutant - *npr^6^* (C-C’), *brZ1* mutant - *rbp^5^* (D-D’), *brZ2* mutant - *br^28^* (E-E’), and *brZ3* mutant - *2Bc^2^* (F-F’). **(G-H)** Quantification of relative regenerative activity using fold change in pouch Dilp8::GFP expression (G) and dorsal hinge Wg expression (H) in late damaged W^1118^ and *br* mutants wing imaginal discs. Fold change determined by normalizing to respective undamaged tissues, ****p<0.0001, **p<0.01, *p<0.05 One-way ANOVA with Tukey’s multiple comparisons tests

To determine whether Broad isoforms participate in the suppression of regenerative activity at the end of larval development, we examined the effect of isoform-specific or pan-isoform disrupting zygotic *br* mutants (in hemizygous males) (Fig. 3B-H) or pan-isoform targeting *br^RNAi^* expression (Fig. S5E) on regenerative signaling. Loss of all Broad isoforms in *npr^6^* or *Bx>br^RNAi^* wing discs allows late-damaged discs to express *dilp8* past the RRP when regenerative activity is usually suppressed (Fig. 3C, G, S5C, F-H). When we examine the induction of regeneration after the RRP using isoform-specific alleles, we determined that BrZ1 and BrZ2 are necessary for restricting *dilp8* expression at the RRP (Fig. 3D, E). Our BrZ3-specific allele *br*(*2Bc^2^*) produced little effect on *dilp8* expression at the regeneration restriction point, consistent with BrZ3 playing a limited role in wing development (Fig. 3F). We were unable to obtain *BrZ4*-specific mutants to examine the loss-of-function phenotypes of this isoform.

When we examine Wg expression in the wing discs of Broad mutants, we observe additional effects of these mutations on both developmental and regenerative Wg expression. In the *npr^6^* mutant, which disrupts all the broad isoforms, we observe a substantial Wg expression reduction in undamaged tissues’ hinge region (Fig. 3C). This reduction is similar to what is seen in the Z2-specific allele *br^28^* (Fig. 3E). In contrast, in *rbp^5^* mutant discs where the Z1 isoform is specifically disrupted, the hinge expression of Wg is largely normal in undamaged discs, but the expression pattern of Wg in the margin is disrupted (Fig. 3D illustrates a representative example). This phenotype may reflect the role of Broad in regulating Cut expression at the margin (Jia et al., 2016), but we leave the further examination of this phenotype for later studies. When we examine tissues damaged after the regeneration restriction point, we see that the *rbp^5^* mutation disabling the Z1 isoform produces a substantial increase in damage-induced Wg expression in the hinge (Fig. S5H). In contrast, the Z2-specific mutation *br^28^* and the pan-isoform mutation *npr^6^* do not produce a significant increase in Wg expression following damage after the regeneration restriction point. We also observe a slight increase in damage-induced Wg expression in Z3-specific mutant *2Bc^2^* (Fig. 3F, H). This increase may reflect a non-autonomous effect of BrZ3 mutation on the regenerating disc. We were unable to evaluate adult tissues to determine whether the increased regenerative signaling activity observed in *br* mutants led to improved tissue repair, as all *br* mutants are either non-pupariating or pupal lethal (D’Avino et al., 1995; Kiss et al., 1988). However, our experiments demonstrate that Broad splice isoforms mediate the ecdysone-dependent restriction of regenerative signaling in late larval wing discs.

### Broad splice isoforms regulate the duration of regenerative activity in damaged discs

Since Broad isoforms are expressed in damaged tissues as the tissues are regenerating (Fig. 3A), we wanted to determine how the loss of Broad or specific Broad isoforms might regulate regenerative activity following early damage of discs (at 80h AED). In early damaged discs, we observe that the pan-isoform mutant *npr^6^* and the Z2 specific mutant *br^28^* produce higher *dilp8* expression12-hours following damage (Fig. S6A-D, E). In addition to differences in the level of *dilp8* expression following damage, we also observe substantial differences in the duration of damage-induced *dilp8* expression between the different mutants, with the pan-isoform mutant *npr^6^* and the Z1-specific mutant, *rbp^5^*, producing *dilp8* expression over a more extended period compared with control discs, or the Z2-specific mutant *br^28^* (Fig. S6A-D, S7A). These results suggest that when damage is produced early when the wing disc can initiate a regenerative response, the duration of *dilp8* expression in the disc is regulated by specific Broad isoforms. We confirmed this result by using *br^RNAi^* to inhibit all the Broad isoforms and demonstrated that *Bx>br^RNAi^* discs also produce extended *dilp8* expression following damage (Fig. S7E-G). In contrast to *dilp8* expression, the effects of the Broad isoform mutants on Wg expression during regeneration are less apparent. As described above, the pan-isoform mutant *npr^6^* and the Z2-specific mutant *br^28^* produce reduced levels of Wg at the hinge region in undamaged tissues (Fig. S6A-D, F S7C). However, all the mutants can produce a similar relative increase in Wg expression following damage (Fig. S7D). Finally, Broad isoforms may also regulate the early events associated with either damage or the initial regenerative response, as we see that the reduction of wing pouch (and overall disc) size following irradiation damage at 80h AED is much greater in all isoform mutants, especially Z1-specific *rbp^5^* and Z2-specific *br^28^* mutant discs (Fig. 6A-D, G, S7B).

In summary, our loss-of-function analysis demonstrates that the individual Broad isoforms play distinct roles in regulating the regenerative signaling response of imaginal discs damaged after the RRP. In addition, the Broad isoforms also regulate the extent and duration of Dilp8 signaling produced by discs damaged before the regeneration restriction point. Based on these observations, we conclude that ecdysone signaling through Broad is necessary to limit both the tissue’s competence to produce a regenerative response, as well as the duration of that response.

### Expression of individual Broad isoforms is sufficient to limit regeneration

Based on our loss-of-function experiments, it appears that the expression of the Broad isoforms may act to limit the duration of regenerative signaling in discs damaged before the RRP or block the initiation of regenerative response in discs damaged after the RRP. To examine whether the expression of individual Broad isoforms is sufficient to limit regeneration, we expressed each of the Broad isoforms in the wing disc and examined both regenerative signaling and the regenerative outcome in adult wings. When we examine regenerative signaling in 80h-damaged discs, we observe that *Bx-Gal4*-driven expression of BrZ1, BrZ2, and BrZ4 limit both *dilp8* and Wg expression in the dorsal compartment of the wing disc, with BrZ1 and BrZ4 producing the strongest inhibition of regenerative Wg expression following damage (Fig. 4A-F, S8A, B). These distinct effects of Broad isoforms on regenerative Wg and *dilp8* expression are also observed in discs experiencing *eiger*-induced damage. BrZ1, BrZ2, and BrZ4 all produce a reduction of *dilp8* expression in these regenerating tissues. However, BrZ1 and BrZ4 produce the strongest inhibition of Wg expression in the *eiger* damage model (Fig. S8 C-I). Therefore, all three isoforms can limit regenerative signaling in *eiger*-damage discs. Consistent with this, RNAi-inhibition of all the Broad isoforms produces elevated levels of both Wg and *dilp8* in *eiger*-damaged tissues (Fig. S8G-I).

**Figure 4.**
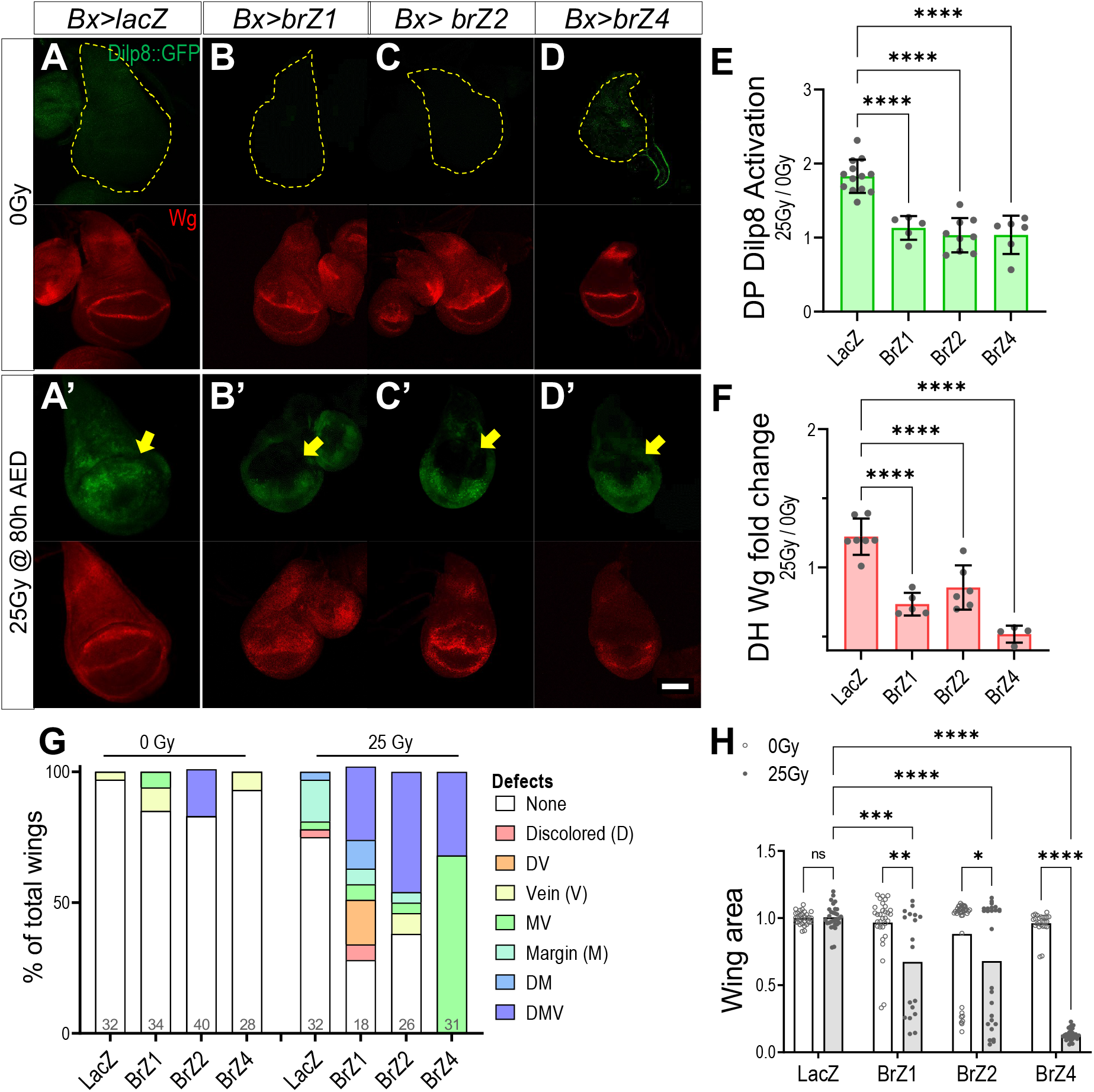
Broad isoform expression is sufficient to suppress regenerative signaling in damaged imaginal discs. **(A-D)** Representative images of pouch Dilp8::GFP (green) and dorsal hinge Wg (red) expression in 92h AED undamaged (A-E), and early damaged - 25Gy @ 80h AED (A’-E’) wing imaginal discs. The yellow dotted line indicates tissue area. Tissues are expressing lacZ (*Bx>lacZ*) as a control (A-A’) or br isoforms, *Bx>brZ1* (B-B’), *Bx>brZ2* (C-C’), and *Bx>brZ4* (D-D’). Primary area of expression indicated by yellow arrows. Scale bar=50um. **(E-F)** Quantification of relative regenerative activity using fold change in dorsal pouch *Dilp8::GFP* expression (E) and dorsal hinge Wg expression (F) in early damaged, control - *Bx>lacZ* and *br* isoform overexpressing wing imaginal discs. Fold change determined by normalizing to respective undamaged tissues, ****p<0.0001, **p<0.01 *p<0.05, One-way ANOVA with Tukey’s multiple comparisons tests. **(G)** Adult wings that show individual defects or combinations of defects increase following late damage. The graph shows the percentage of defective adult wings from larvae transiently overexpressing *lacZ, brZ1, brZ2*, and *brZ4 (rn-gal4*) with no damage (0Gy) and early damage (25Gy) during wing development. Population size is indicated in the graph. **(H)** Quantification of adult wing size for tissue in (G). Size of wing measured in unit area and normalized to undamaged *rn>lacZ* wing size of respective sex. Two-way ANOVA with Tukey’s multiple comparisons test, ****p<0.0001

Constitutive expression of individual Broad isoforms produces substantially deformed adult wings, making it challenging to assess regenerative outcomes. Therefore, to determine whether expression of the individual Broad isoforms is sufficient to inhibit regeneration, we transiently expressed each of the Broad isoforms in the wing pouch using *rn-Gal4* and used *tub-Gal80^ts^* expression to limit the timing of expression to a 12-hour window following irradiation (Fig. S9A). We observe that transient expression of BrZ1, BrZ2, and BrZ4 in undamaged control larvae produces only minor effects on disc patterning and growth (Fig. 4GH, S9B-E). However, the transient expression of Broad isoforms following early third instar X-irradiation damage profoundly affects wing regeneration. Expression of BrZ1, BrZ2, and BrZ4 results in a high proportion of incompletely regenerated discs and reduced wing blade size. Of the three splice isoforms, BrZ4 produces the most potent inhibition of regeneration, with all the adult wings mis-patterned and extremely small (Fig. 4G, H, S9B-E).

In summary, our isoform expression experiments demonstrate that the local expression of individual Broad isoforms in damaged tissues is sufficient to block critical local and systemic regeneration signaling events. Even the transient expression of single Broad isoforms in regenerating tissues can severely attenuate regeneration in these tissues.

### Ecdysone inhibits and promotes *wg* expression through distinct pathways

We have demonstrated here that ecdysone produces dual effects on regenerative signaling. At lower levels, ecdysone can promote the expression of Wg and *dilp8* in regenerating tissues. At higher levels, ecdysone limits regeneration and the expression of these regenerative signals through the activation of Broad splice isoforms. To better understand how ecdysone produces these distinct effects on regenerative signaling, we examined how ecdysone regulates a regulatory region located ~8 kb downstream of the *wg* coding region (Fig. S10A), which was previously named BRV118 (Schubiger et al., 2010) but has more recently been described as the *wg* Damage Responsive Enhancer (*wg*DRE; (Harris et al., 2020)). The *wg*DRE is critical for the regenerative activation of *wg* expression following damage. Epigenetic changes at the *wg*DRE towards the end of larval development lead to the loss of regenerative capacity following the RRP (Harris et al., 2016). Using a transgenic reporter of *wg*DRE activity (*wg*DRE-GFP, (Harris et al., 2016), Fig. S10A), we observe the attenuation of damage-induced *wg*DRE activity as larvae develop past the RRP (Fig. S10B, C). To first determine whether the limitation of regenerative activity by ecdysone following the regeneration restriction point is mediated through the *wg*DRE, we measured reporter expression in *Bx>EcR.A^DN^* discs. We see that blocking ecdysone signaling increases the damage-induced wgDRE reporter activity in the dorsal pouch of discs damaged after the RRP (Fig. 5A, B, D. S10D). Therefore, the inhibition of the *wg*DRE in late-damaged tissues is dependent on ecdysone signaling in regenerating disc tissues. Consistent with our earlier observations, we also see that the inhibition of the *wg*DRE after the regeneration restriction point is dependent on Broad, as late damage can also activate the *wg*DRE in *Bx>br^RNAi^* expressing discs (Fig 5A,C,D, S10D). To determine whether Broad isoform expression is sufficient to suppress *wg*DRE activity, we examined whether the damage-induced activation of *wg*DRE before the regeneration restriction point can be suppressed by expression of Broad isoforms. We see that expression of BrZ1, BrZ2, and BrZ4 can suppress expression of the *wg*DRE following early damage (Fig. 5E-I, S10E). Based on these results, we conclude ecdysone, via Broad isoform expression, can limit regenerative activity by suppressing the damage-induced activity of the *wg*DRE.

**Figure 5.**
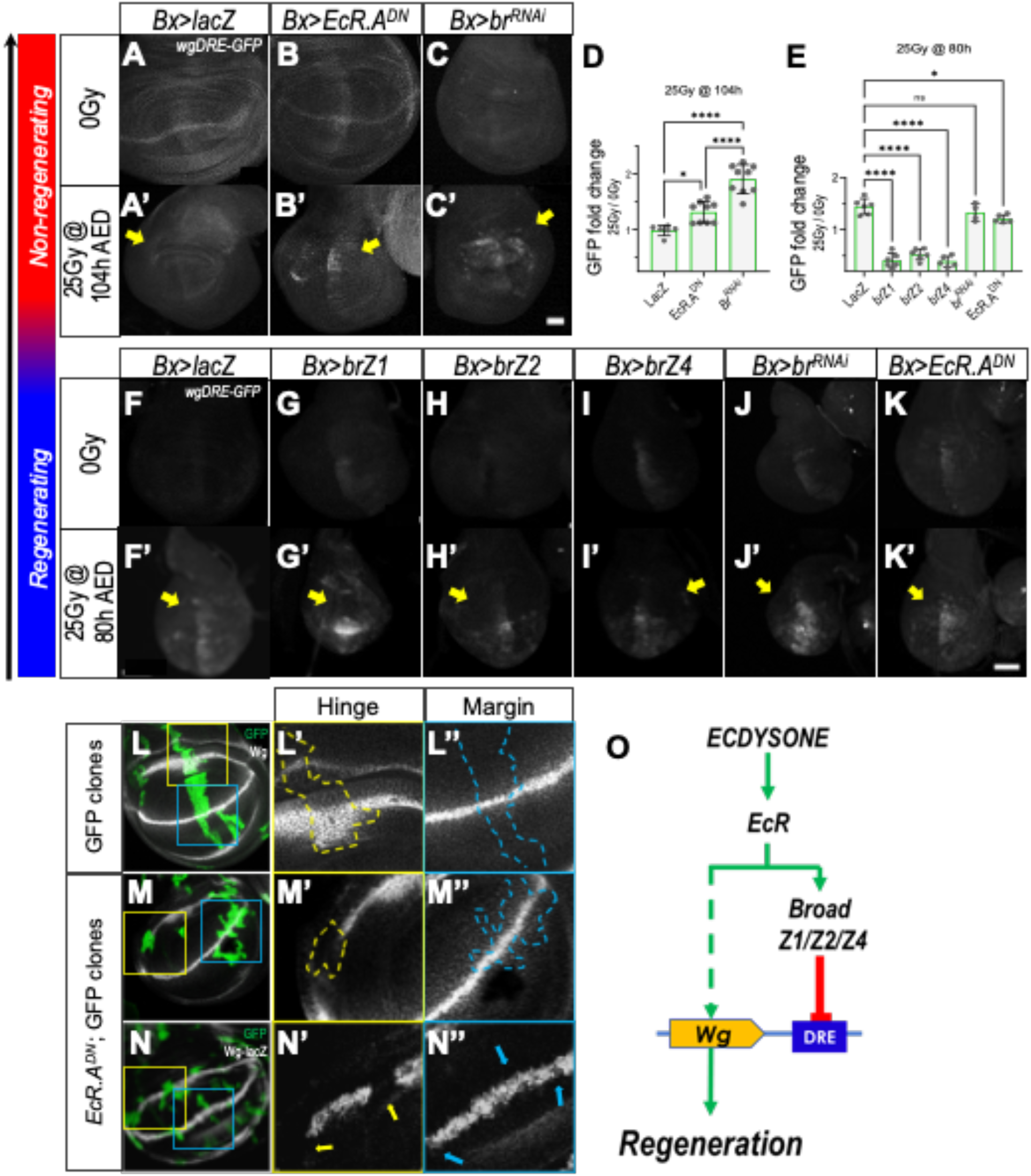
Ecdysone signaling regulates Wg expression. **(A-C)** Representative images of wgDRE-GFP (grey) expression in 116h AED undamaged (A-C) and late damaged - 25Gy @ 104h AED (A’-C’) wing imaginal discs. Yellow arrows indicate the primary area of Gal4-UAS expression. Tissues are expressing lacZ (*Bx>lacZ*) as a control (A-A’), *Bx>EcR^DN^* (B-B’), and *Bx>br^RNAi^* (C-C’). Scale bar=50um. **(D)** Quantification of dorsal pouch wgDRE-GFP expression fold change following late damaged (25Gy-104h), control - *Bx>lacZ, Bx>EcR^DN^* and *Bx>br^RNAi^* overexpressing wing imaginal discs. Quantification normalized to respective undamaged tissues. ****p<0.0001, *p<0.05, One-way ANOVA with Tukey’s multiple comparisons tests. **(E)** Quantification of dorsal pouch *wg*DRE-GFP expression fold change following early damaged, control - *bx>lacZ, br* isoform overexpressing, *br* knockdown, and *EcR^DN^* expressing wing imaginal discs. Normalized to respective undamaged tissues, ****p<0.0001, **p<0.01, One-way ANOVA with Tukey’s multiple comparisons test. **(F-K)** Representative images of *wg*DRE-GFP (grey) expression in 92h AED undamaged (F-K) and early damaged - 25Gy @ 80h AED (F’-K’) wing imaginal discs. Tissues are expressing lacZ (*Bx>lacZ*) as a control (F-F’) or br isoforms, *Bx>brZ1* (G-G’), *Bx>brZ2* (H-H’), *Bx>brZ4* (I-I’), *Bx>br^RNAi^* (J-J’), and *bx> EcR.A^DN^* (K-K’). Primary area of expression indicated by yellow arrows. Scale bar=50um. **(L-M)** Representative images of Wg (grey) expression in control, *UAS-GFP* alone (L), and *UAS-EcR.A^DN^;UAS-GFP* (M), MARCM clones. Larvae were heat-shocked at 60h AED, and tissues were isolated at 104h AED. Zoom-in images of clones at the hinge (L’-L’’) and margin (M’-M’’) are shown on the right. Arrows indicate the region where clones cross the Wg expression at the hinge (yellow) and margin (blue). **(N)** Wg locus transcriptional activity - *UAS-EcR.A^DN^;UAS-GFP* clones in *wg*-lacZ background. Clones at hinge and margin are shown in N’ and N’’ respectively. **(O)** Ecdysone signaling cascade for the regulation of Wg expression.

To assess how lower levels of ecdysone function to promote regenerative Wg expression, we first examined how the loss of ecdysone signaling affects Wg expression in undamaged wing discs. We had observed in Figure 1D, *Bx>EcR.A^DN^* expression appears to suppress hinge Wg expression in undamaged tissues. To examine this more carefully, we used MARCM to generate GFP-labeled clones that expressed *EcR.A^DN^*. We observed the expression of *EcR.A^DN^* produced clones that cell-autonomously inhibited Wg expression at the hinge regions of the developing wing disc but not at the margin (Fig. 5L, M). This inhibition appears to be a transcriptional regulation of *wg* expression as we see a similar effect of *EcR.A^DN^* expression on the activity of a *wg* transcriptional reporter line (*wg-lacZ*, Fig. 5N). It is possible that ecdysone exerts both its inhibitory and activating effects on Wg expression through the *wg*DRE. However, when we inhibit ecdysone signaling (*Bx>EcR.A^DN^*) or Broad isoform expression (*Bx>br^RNAi^*) in early-damaged discs, we see that neither of these manipulations limit *wg*DRE activation (Fig. 5E, J, K, S10E). Therefore, ecdysone signaling is required for regenerative Wg expression but regulates *wg* transcription independently of Broad and through a regulatory region that is not part of the *wg*DRE. These distinct pathways for ecdysone regulation of Wg are summarized in Figure 5O.

### Ecdysone signaling coordinates regeneration with the duration of the larval period

Damage and regeneration of imaginal discs activates a regeneration checkpoint, a delay in development that extends the larval period. This checkpoint arises from the expression and release of Dilp8 from regenerating tissues (Colombani et al., 2012; Garelli et al., 2012). Dilp8 binds to its receptor Lgr3, expressed in both the brain and the ecdysone-producing prothoracic gland, to limit the ecdysone synthesis (Colombani et al., 2015; Garelli et al., 2015; Jaszczak et al., 2016). This Dilp8-Lgr3 signaling delays the accumulation of ecdysone at the end of the larval period, extending the regenerative competence of imaginal discs and delaying the transition to the pupal phase of development (Garelli et al., 2015; Jaszczak et al., 2016). Because Dilp8 activation of the regeneration checkpoint extended the regenerative period of development, we (Jaszczak et al., 2015, 2016; Jaszczak & Halme, 2016) and other researchers (Andersen et al., 2013; Colombani et al., 2012, 2015; Garelli et al., 2012; Vallejo et al., 2015) have hypothesized that the extra time was required for the additional growth and repatterning required to complete regeneration. However, since we have shown that ecdysone produces a biphasic effect on regenerative signals in the damaged disc, we wanted to test the hypothesis that checkpoint delay was required to accommodate regeneration. To do this, we examined regeneration in homozygous *dilp8^-^* larvae, which produce minimal checkpoint delay following damage (Fig. 6A), and therefore have a shorter regenerative period. Unexpectedly, adult flies arising from X-irradiated *dilp8^-^* larvae could regenerate their wing discs as successfully as control *dilp8*^+^ (*w^1118^*) adults. The lack of regeneration checkpoint delay produces no significant impact on either tissue repatterning (Fig. 6B) or regrowth to target undamaged tissue size. In fact, we see that *dilp8^-^* larva produce regenerated adult wings that are closer to the undamaged target size than *dilp8^+^* larvae in which the checkpoint is intact. (Fig. 6C, S11A).

**Figure 6.**
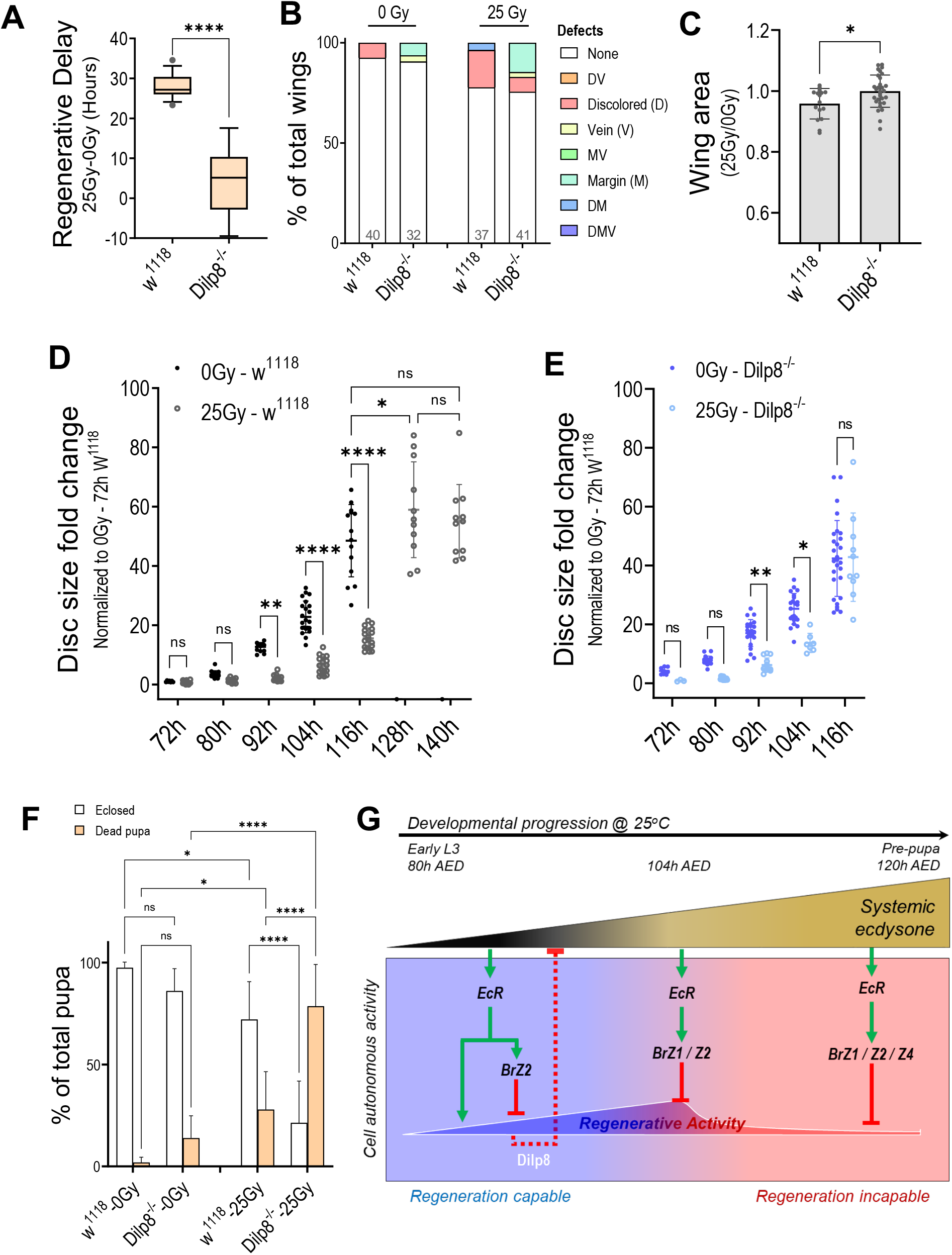
*Dilp8^-^* larvae meet regeneration targets within the attenuated development period. **(A)** Quantification of regenerative delay following early damage (20Gy-80h) in *w^1118^ (dilp8^+^*) and *dilp8^-/-^* larvae. Unpaired t-test, ****p<0.0001. **(B)** Adult wings that show individual defects or combinations of defects increase following late damage. The graph shows the percentage of defective adult wings following no damage (0Gy) and early damage (25Gy-80h) in *w^1118^* and *dilp8^-/-^* adults. Population size is indicated in the graph. **(C)** Quantification of adult wing size for tissue in (B). Size of wing measured in unit area and normalized to undamaged wing size of respective genotype and sex. Unpaired t-test, *p<0.05. **(D-E)** Quantification of wing disc growth following no damaged tissues (0Gy) and damage (25Gy) at 48h AED in *w^1118^* (D) and *dilp8^-/-^* (E) larvae. Data sets normalized to 72h undamaged *w^1118^* tissues. ****p<0.0001, ***p<0.001, **p<0.01, *p<0.05, 2-way ANOVA with Tukey test. **(F)** Quantification of population viability following early damage (20Gy-80h) in *w^1118^* and *dilp8^-/-^* larvae. ****p<0.0001, *p<0.05, 2-way ANOVA with Tukey test. **(G)** The summary model illustrates the sequence of ecdysone-regulated cell-autonomous events that regulate regenerative activity and regenerative capacity during wing disc development.

To better understand how *dilp8^-^* larvae can regenerate their damaged wing discs despite the attenuated regenerative period, we measured disc size in undamaged and regenerating discs through the third instar of *dilp8^+^* and *dilp8^-^* larvae. In control larvae, damage produces a delay in disc growth, with regenerating imaginal discs being measurably smaller than undamaged controls between 92 and 116h AED, just before unirradiated larvae typically end their larval period. However, during the extended larval period produced by activation of the regenerative checkpoint, the growth of the regenerating imaginal discs rapidly reaches the target size (the final size of the undamaged discs) by 128h AED and then remain at that size until the end of the larval period (Fig. 6D). In *dilp8^-^* larvae, we still observe a growth lag in damaged and regenerating tissues, with regenerating tissues being significantly smaller between 92 and 104h AED. However, unlike *dilp8^+^* larvae, the regenerating imaginal discs in *dilp8^-^* larvae rapidly grow after 104h AED, reaching target size by 116hAED, just before both control and *dilp8^-^* larvae pupate (Fig. 6E). When we directly compare the growth of control and *dilp8^-^* imaginal discs, we see that undamaged discs grow at approximately the same rate (Fig. S11B), whereas regenerating *dilp8^-^* imaginal discs grow much faster than control discs (Fig. S11C). In summary, we observe that in the absence of the regeneration checkpoint, ecdysone synthesis is no longer limited, and the regenerative growth of imaginal discs is accelerated such that target disc size is still reached by the end of the shortened larval period. This suggests that the biphasic effect of ecdysone signaling on disc regeneration is capable of coordinating disc regeneration with the duration of the larval period.

While Dilp8 and checkpoint activation are not necessary for providing additional time to accommodate regenerative growth, checkpoint activity is essential for maintaining the viability of pupae following regeneration. The frequency of pupal lethality (pupal cases where the adults fail to eclose) in control larvae is relatively low, typically ~25% for larvae irradiated at 25Gy. However, pupal lethality of irradiated dilp8^-^ larvae is much higher, ~78% (Fig. 6F). We believe that the increase in pupal lethality is a consequence of regenerative activity in *dilp8^-^* larvae as control larvae irradiated late (post-RRP), which fail to initiate a regenerative response, still produce a relatively low rate of pupal lethality (~30%, Fig. S11D). Therefore, Dilp8 checkpoint activation appears to play an essential role in preserving the future pupal viability of animals undergoing disc regeneration.

## Discussion

Following damage, regenerating *Drosophila* imaginal discs release Dilp8, which circulates in the larval hemolymph and signals through its receptor Lgr3 in the larval brain and the PG to limit ecdysone production (Colombani et al., 2012, 2015; Garelli et al., 2012; Jaszczak et al., 2016; Vallejo et al., 2015). By delaying the increase of ecdysone that signals the end of larval development, Dilp8 extends the regenerative period (Colombani et al., 2012; Garelli et al., 2012). However, we demonstrate here that even in the absence of Dilp8 signaling, damaged wing imaginal discs are capable of the repatterning and growth required to reach their regeneration target in an attenuated regenerative period (Fig. 6B, C). However, we see that the accelerated regeneration in *dilp8^-^* larvae is also accompanied by a substantial increase in pupal lethality (Fig. 6F). This pupal lethality does not appear to result from unrepaired damage, as larvae irradiated after the RRP when they cannot initiate a regenerative response do not show elevated levels of pupal lethality (Fig. S11D). Therefore, the role of Dilp8 and regenerative checkpoint signaling may be primarily to preserve viability in the presence of regenerating tissues as opposed to providing adequate time for regeneration. In this study, we did not examine how regeneration checkpoint activation preserves pupal viability. Still, one possibility is that an extended larval feeding period may allow larvae with regenerating tissues the ability to store up sufficient energy reserves to both regenerate damaged tissues and complete metamorphosis. Further study is necessary to understand better how regeneration impacts pupal viability.

In contrast to *dilp8^-^* larvae, which can produce completely regenerated tissues in an attenuated regenerative period, artificially increasing ecdysone levels through feeding increases the number of incompletely regenerated tissues substantially (Fig. S2C, D). This suggests that the changing ecdysone levels that occur throughout the regenerative period of larval development exert sensitive control over imaginal disc regeneration and that this regulation is lost when we ectopically manipulate ecdysone levels. In this study, we demonstrate that ecdysone plays a dual role in regulating regenerative activity in damaged discs to ensure that regeneration targets are achieved within the larval period. (Summarized in Fig. 6G). First, we demonstrate that ecdysone signaling in damaged imaginal discs is necessary for the upregulation of Wg and Dilp8, critical signals mediating the local and systemic regenerative responses (Fig. 1L, M, N). This observation may reflect a requirement for ecdysone signaling in Wg expression at the hinge region (Fig. 5L, M, N), which is critical to irradiation-induced regenerative responses (Verghese & Su, 2016). As development progresses through the third instar, ecdysone pulses produced by the prothoracic gland (PG) cause ecdysone to accumulate in the larvae (Lavrynenko et al., 2015). We propose that this accumulation initiates a series of concentration-dependent events (Fig. 2B-F). First, ecdysone promotes regeneration, accelerating regenerative activity as ecdysone titer increases. Then, when ecdysone titer reaches a high level in the larvae, ecdysone signaling suppresses regenerative activity by activating the expression of the Broad splice isoforms (Fig 3), which suppress regenerative activity in damaged discs (Fig. 4).

This biphasic, concentration-dependent regulation of regenerative activity allows ecdysone to coordinate regeneration with the duration of larval development. This coordinating role may be similar to how ecdysone coordinates imaginal disc patterning with the larval development period (Alves et al., 2020). Similarly, recently published experiments have observed that damage of the *Drosophila* hindgut during L2 or early L3 produces Dilp8, delaying the onset of pupariation, whereas damage to the hindgut of wandering L3 larvae no longer produces Dilp8 or developmental delay. However, regeneration is still completed in this attenuated period through accelerated mitotic cycling, which allows the tissue to meet the regeneration target within the mitotic regeneration window (Cohen et al., 2021). In this example, while ecdysone signaling regulates the end of mitotic regeneration, the role of ecdysone signaling in producing the accelerated mitoses has not been investigated.

Other pathways have been implicated in regulating regenerative activity in the wing disc or the loss of regenerative capacity at the end of development. Further experiments will be required to determine how these pathways are regulated by ecdysone and Broad. Recent work has demonstrated that Chinmo, which regulates the ‘stemness’ of cells, is expressed during early wing disc development and antagonizes BrZ1 to allow regenerative activity (Narbonne-Reveau & Maurange, 2019). While ecdysone signaling suppresses Chinmo expression at the RRP (Narbonne-Reveau & Maurange, 2019), Chinmo has also been shown to regulate EcR activity in other tissues (Marchetti & Tavosanis, 2017). It is unclear whether ecdysone and Chinmo interact during early larval development. At the end of larval development, the *Drosophila* genome undergoes extensive epigenetic changes in preparation for pupation and metamorphosis (Ma & Buttitta, 2017; Saha et al., 2019). The Polycomb-group proteins (PcG) produce silencing modifications on the heterochromatin at *wg*DRE to block regeneration after the RRP (Harris et al., 2016). However, it remains unclear how PcG is recruited to the wgDRE. There is a putative Pc-binding site in the *wg*DRE, but it is not necessary for silencing this locus at the end of larval development (Harris et al., 2016). Both PcG and Broad are essential for suppressing regeneration genes, and there is evidence that Broad and PcG physically interact during the development of the wing disc (Lv et al., 2016). Therefore, it is possible that Broad isoforms may recruit PcG to wgDRE to suppress regeneration at the end of larval development.

In summary, our findings provide new insight into the mechanisms that coordinate tissue regeneration with the development of the animal as a whole. The steroid hormone ecdysone has a biphasic, concentration-dependent effect on the regenerative activity of the *Drosophila* wing imaginal disc. Through this biphasic signaling, ecdysone can coordinate the completion of regeneration with the end of the larval period of growth in wild-type larvae and larvae that lack the regeneration checkpoint, where the regenerative period is attenuated. We demonstrate that the regeneration checkpoint may be important for maintaining pupal viability in animals that have regenerated their imaginal discs.

## Materials and Methods

### Drosophila stocks and culture

Stocks used include w^1118^ (BDSC_5905), Bx-Gal4;UAS-Dcr2; (Bilder lab stock), Dilp8::GFP/TM6B (Derived from BDSC_33079), UAS-LacZ.NZ (BDSC_3956), UAS-EcR.A^W650A^ (BDSC_9451), y^1^,br^2Bc-2^/Binsn (BDSC_29969), y^1^,br^npr-6^/Binsn (BDSC_36562), y^1^,br^rbp-5^/Binsn (BDSC_30138), y^1^,br^28^,w^1^/Binsn (BDSC_36565), FM7c^tb^ (BDSC_36337), UAS-BrZ1 (BDSC_51190), UAS-BrZ2 (BDSC_51191), UAS-BrZ3 (BDSC_51192), UAS-BrZ4 (BDSC_51193), UAS-Dcr2;UAS-BrRNAi (Derived from BDSC_27272), Bx-Gal4;UAS-Eiger; (Derived from regg^1^ stock, from H. Kanda), wg^1^,FRT40A;Dilp8::GFP/SM6-TM6B (Derived from BDSC_2978 and BDSC_33079), Bx-Gal4;wgDRE-GFP;dcr (Derived from Hariharan lab BRV118-GFP stock), UAS-mCD8-GFPhsFlp;tub-Gal4;FRT82B,tubGal80/TM6B (Siegrist lab stock), UAS-EcR.A[W650A]; FRT82B/SM6-TM6B (Derived from BDSC_9451), Ubx-Flp;FRT40A;FRT82B (BDSC_42733), wg-LacZ,UAS-EcR.A[W650A];FRT82B/CyO (Derived from BDSC_11205 and BDSC_9451)

Experimental lines and crosses were maintained at 25°C with a 12-hour alternating light-dark cycle. The developmental timing was synchronized by staging egg-laying on grape agar plates (Genesee Scientific) during a designated 4-hour interval. Twenty-four hours after egg deposition (AED), 20 first-instar larvae were transferred into vials or plates containing standard (cornmeal-yeast-molasses) media (Archon Scientific B101). The larvae remained undisturbed in the media at 25°C or 18°C until treatments began at the third larval instar.

### Irradiation Damage and Ecdysone Feeding

Staged larvae were either left undamaged or exposed to 20 or 25 Gy X-irradiation generated from a 43805N X-ray system Faxitron operating at 130 kV and 3.0 mA. The larvae were exposed to x-irradiation at 80h AED for early damage, 104hAED for late damage, and 92hAED in experiments in S.1 & S.10.

Ecdysone food was prepared by dissolving 20-hydroxyecdysone (Sigma – starting concentration: 20mg/ml in 95% ethanol) in 2 ml of food media at final concentrations of 0.1, 0.3, 0.6 and 1.0 mg/ml of food, or an equivalent volume of 95% ethanol (0 mg/ml) for control. Larvae were reared as previously described until 80h AED then transferred to the ecdysone or ethanol-control food, approximately 6-7 larvae per vial (Halme et al., 2010).

### Pupariation Time and Developmental Delay

For calculating purposes, 0h AED was considered to be the middle of the egg-laying interval. The pupae in each vial were counted approximately every 12 hours, starting around 104h AED and ending three days after the most recent pupation. The data were pooled from multiple vials of the same genotype laid on the same day. Data from separate lays were calculated separately, and at least three lays are represented in each experiment. Median pupariation time was then calculated (Equation 1). Developmental delay was considered to be the difference in pupariation time between the experimental and control groups.

***Equation 1:** Median pupariation time calculation*

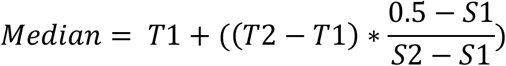

Median pupariation time was calculated by first determining the sum fraction of total pupae counted at each time point for each genotype. The first time point with a sum fraction of total pupae exceeding 50% indicates that the median pupariation time occurred between that point and the proceeding time point. We next calculated how long past the proceeding timepoint 50% of larvae pupated and the difference between the sum fractions. To determine how far past the first timepoint the median pupariation time was, we divided the difference from the halfway point by the difference between the sum fractions then multiplied this by the difference between the time points. We then added this number to the preceding time point. T2 indicates the later timepoint, T1 indicates the earlier timepoint, S2 indicates the sum fraction of pupae at T2, S1 indicates the sum fraction of pupae at T1.

### Tissue Isolation

#### Adult wings

adult flies were approximately 36 hours after eclosion, separated according to sex, and stored in 70% ethanol. No staining or tissue treatment was required. Wings were isolated and mounted onto slides using Gary’s Magic Mounting (GMM – Balsam powder dissolved in methyl salicylate) media.

#### Wing discs for imaging

Larvae were inverted and cleaned in PBS. The larvae carcass (cuticle with attached imaginal discs) was then fixed with 4% paraformaldehyde in PBS for 20 mins, followed by two 5-min washes in PBS. In broad mutant experiments, only the imaginal discs of hemizygous male larvae were isolated. To distinguish male from female larvae, we visually identified the gonads found in the lower abdominal flank of male larvae only. Male larvae gonads appear as circular translucent discs visible through the cuticle (Selva & Stronach, 2007).

#### Wing discs for Western Blot

40-80 imaginal discs (depending on tissue size at each time point) were isolated from third instar larvae. All tissues were isolated in chilled Schneider’s Insect Medium (Sigma-Aldrich). The dissection dish was placed on ice during the entire dissection. Isolated tissues were washed twice in chilled PBS and spun down for 15 seconds using a C1008-R Benchmark myFUGE mini centrifuge. Excess PBS was aspirated out, then the tissues (in approximately 50ul of PBS) were frozen in dry ice before storage at −80°C.

### Immunofluorescent Staining

The isolated imaginal discs were permeabilized for immunofluorescent staining using two 10-min washes in 0.3% Triton in PBS (PBST), then incubated for 30 mins in a blocking solution of 10% goat serum (GS) and 0.1% PBST. Then the tissues were incubated in primary antibody solutions overnight at 4°C on a nutator. Antibody solutions were prepared in 10% GS in 0.1% PBST. The primary antibodies used are mouse β-Gal (1:250; Promega #Z3781), mouse anti-Wingless (1:100; DSHB #4D4), and rabbit anti-GFP (1:1000; Torrey Pines Biolabs #TP401). The samples were washed and blocked again before incubating in the appropriate secondary antibody solutions (1:1000; ThermoFisher Alexa488, Cy3, or Alexa633) prepared in 10% GS in 0.1% PBST for 2-4 hours at room temperature (RT). After two 10-min 0.3% PBST washes and one 5-min PBS wash, the tissues were stored in 80% glycerol in PBS at 4°C. The tissues were mounted for imaging within a week of staining. During mounting, imaginal discs were isolated from the stained carcass and mounted on glass slides with Vectashield (Vector Laboratories).

### Imaging, Quantification and Statistical Analysis

Adult wings were imaged using MU530-Bi AmScope Microscope Digital Camera and software. Confocal imaging was done using an Olympus FluoView 1000 from the University of Virginia Department of Cell Biology and Zeiss LSM 700 and LSM 710 in the University of Virginia Advanced Microscopy Facility (RRID: SCR_018736). Laser power and gain settings for each set of stained samples were based on the experimental group with the highest fluorescence intensity in each channel and kept constant within the experiment. All images were taken as z-stacks of 10um intervals. Images were processed and quantified with Fiji/ImageJ. Representative images used in figures are composites of the image stacks using max fluorescence projection, while quantification was done using sum fluorescence composites.

The Wingless and GFP (Dilp8 and wgDRE) quantification region was determined differently in undamaged and irradiation damaged tissues versus tissues damaged by eiger expression (described below). In undamaged and irradiated wing imaginal discs, Wg was quantified in the dorsal hinge of the imaginal disc pouch by tracing Wingless in this region from the dorsal edge of the margin. Margin Wg was not quantified, as is it not associated with the regenerative activity. Ventral hinge Wingless was not quantified as tissue evagination or folding during mounting often interfered with distinguishing margin and ventral hinge Wg. GFP fluorescence was quantified in the pouch region of the discs as defined by the outer edge of hinge Wg expression surrounding the wing pouch. In eiger damaged tissues, the blastema area was determined by the area of GFP (Dilp8) expression, and only the Wg within the area of GFP expression was quantified. In quantification of each damage model, the expression of Wg or GFP in the notum was not quantified. However, quantification of wing imaginal disc size (unit area) included both wing pouch and notum.

Prism 8 software was used for Statistical Analysis. To compare between independently repeated experiments, we normalized within the experiment as indicated in figure legends. The specific tests that were used are listed in the figure descriptions.

### Western Blot

Proteins were extracted in 50ul SDS lysis buffer (2% SDS, 60mM Tris-Cl pH6.8, 1X protease inhibitors, 5mM NaF, 1mM Na orthovanadate, 1mM β glycerophosphate in distilled H_2_O), sonicated using two 5-sec pulses (microtip Branson sonifier), boiled for 10 minutes at 95°C and centrifuged at 15000rpm for 5 mins at RT. The supernatant was collected for BCA assay and analyzed by SDS-PAGE using Mini-Protean® TGXTM 4– 15% (BioRad) and transferred to nitrocellulose membranes. For Western blot analysis, membranes were incubated with blocking solution (1% cold water fish gelatin; Sigma #G7765), primary antibodies (1:500 Broad Core, DSHB #25E9.D7 and 1:10,000 a-tubulin, Sigma #T6074), followed by appropriate LI-COR IRDye® secondary antibodies and visualized using the Li-COR Odyssey® CLx Imaging System. Quantifications were calculated with LI-COR Image Studio™ Software.

## Acknowledgments

The authors would like to thank Iswar Hariharan and Rob Harris for *wg*DRE reporter lines and Andre Landin Malt and Brittany Martinez for WB reagents and assistance. The authors would like to acknowledge the University of Virginia Advanced Microscopy Facility (RRID: SCR_018736) for training and access to the Zeiss LSM700 and LSM710 confocal microscopes used in this study.

## Data Availability

All data underlying this work will be made available upon request.

