## Supplemental data and Figures for "Ecdysone exerts biphasic control of regenerative signaling, coordinating the completion of regeneration with developmental progression"

### Supplementary Materials and Methods

#### Genotypes

Figure 1

A-D: $w^{1118}$

E: Dilp8$^{MI00727}$ /+

F: $w^{1118}$

G, K: Bx-Gal4 / +; UAS-Dcr2 / +; Dilp8$^{MI00727}$ / UAS-LacZ.NZ

H, L: Bx-Gal4 / +; UAS-Dcr2 / UAS-EcR.A$^{W650A}$; Dilp8$^{MI00727}$ /+

I-J: genotypes in Fig.1 G-H

M-N: genotypes in Fig.1 K-L

Figure 2

B-I: Dilp8$^{MI00727}$ / +

Figure 3

A-B: $w^{1118}$

C: br$^{npr-6}$; +/+; Dilp8$^{MI00727}$/+

D: br$^{rbp-5}$; +/+; Dilp8$^{MI00727}$/+

E: br$^{28}$; +/+; Dilp8$^{MI00727}$/+

F: br$^{2Bc-2}$; +/+; Dilp8$^{MI00727}$/+

G-H: genotypes in Fig.2 B-F

Figure 4

A: Bx-Gal4 / +; UAS-Dcr2 / +; Dilp8$^{MI00727}$ / UAS-LacZ.NZ

B: Bx-Gal4 / +; UAS-Dcr2 / UAS-BrZ1; Dilp8$^{MI00727}$ / +

C: Bx-Gal4 / +; UAS-Dcr2 / UAS-BrZ2; Dilp8$^{MI00727}$ / +

D: Bx-Gal4 / +; UAS-Dcr2 / UAS-BrZ4; Dilp8$^{MI00727}$ / +

E-F: genotypes in Fig.4 A-D

G-H: AP1-GFP/ +; rn-Gal4, tub-Gal80$^{ts}$ / UAS-LacZ.NZ

AP1-GFP / UAS-BrZ1; rn-Gal4, tub-Gal80$^{ts}$ / +

AP1-GFP / UAS-BrZ2; rn-Gal4, tub-Gal80$^{ts}$ / +

AP1-GFP / UAS-BrZ4; rn-Gal4, tub-Gal80$^{ts}$ / +

Figure 5

A, F: Bx-Gal4 / +; BRV118-GFP / +; UAS-Dcr2 / UAS-LacZ.NZ

B, K: Bx-Gal4 / +; BRV118-GFP / UAS-EcR.A$^{W650A}$; UAS-Dcr2 / +

C, J: Bx-Gal4 / +; BRV118-GFP / UAS-Br$^{RNAi}$; UAS-Dcr2 / +

G: Bx-Gal4 / +; BRV118-GFP / UAS-BrZ1; UAS-Dcr2 / +

H: Bx-Gal4 / +; BRV118-GFP / UAS-BrZ2; UAS-Dcr2 / +

I: Bx-Gal4 / +; BRV118-GFP / UAS- BrZ4; UAS-Dcr2 / +

L: UAS-mCD8-GFP, hs-Flp; tub-Gal4 / +; FRT82B, tub-Gal80 / FRT82B

M: UAS-mCD8-GFP, hs-Flp; tub-Gal4 / UAS-EcR.A$^{W650A}$; FRT82B, tub-Gal80 / FRT82B

N: UAS-mCD8-GFP, hs-Flp; tub-Gal4 / wg$^{en-11}$, UAS-EcR.A$^{W650A}$; FRT82B, tub-Gal80 / FRT82B

Figure 6

A-C, F: w$^{1118}$ and Dilp8$^{MI00727}$ / Dilp8$^{MI00727}$

D: w$^{1118}$

E: Dilp8$^{MI00727}$ / Dilp8$^{MI00727}$

Figure S1

A: w$^{1118}$

B-C: Dilp8$^{MI00727}$ /+

D-E: w$^{1118}$

Figure S2

A-H: w$^{1118}$

Figure S3

B-E: Bx-Gal4 / +; UAS-Dcr2 / +; Dilp8$^{MI00727}$ / UAS-LacZ.NZ

Bx-Gal4 / +; UAS-Dcr2 / UAS-EcR.A$^{W650A}$; Dilp8$^{MI00727}$ / +

F: Bx-Gal4 / +; UAS-Eiger / +; Dilp8$^{MI00727}$ / UAS-LacZ.NZ

F': Bx-Gal4 / +; UAS-Eiger / UAS-EcR.A$^{W650A}$; Dilp8$^{MI00727}$ / +

G-H: genotypes in Fig.S3 F-F'

Figure S4

A-C: w$^{1118}$

D-F: Bx-Gal4 / +; UAS-Eiger / +; Dilp8$^{MI00727}$ / +

Figure S5

B: w$^{1118}$

C-D: w$^{1118}$

br$^{npr\text{-}6}$; +/+; Dilp8$^{MI00727}$/+

br$^{rbp\text{-}5}$; +/+; Dilp8$^{MI00727}$/+

br$^{28}$; +/+; Dilp8$^{MI00727}$/+

br$^{2Bc\text{-}2}$; +/+; Dilp8$^{MI00727}$/+

E: Bx-Gal4 / +; UAS-Dcr2 / UAS-Br$^{RNAi}$

F: Bx-Gal4 / +; UAS-Dcr2 / +; Dilp8$^{MI00727}$ / UAS-LacZ.NZ

G: Bx-Gal4 / +; UAS-Dcr2 / UAS-Br$^{RNAi}$; Dilp8$^{MI00727}$ / +

H: genotypes in Fig.S5 F and G

Figure S6

A: w$^{1118}$

B: $br^{npr-6}$; +/+; $Dilp8^{MI00727}$/+

C: $br^{rbp-5}$; +/+; $Dilp8^{MI00727}$/+

D: $br^{28}$; +/+; $Dilp8^{MI00727}$/+

E-G: genotypes in Fig.S6 A-D

Figure S7

A-D: Genotypes in S6 above

E, G: Bx-Gal4 / +; UAS-Dcr2 / +; $Dilp8^{MI00727}$ / UAS-LacZ.NZ

F, G: Bx-Gal4 / +; UAS-Dcr2 / UAS-$Br^{RNAi}$; $Dilp8^{MI00727}$ / +

Figure S8

A-B: Bx-Gal4 / +; UAS-Dcr2 / +; $Dilp8^{MI00727}$ / UAS-LacZ.NZ

Bx-Gal4 / +; UAS-Dcr2 / UAS-BrZ1; $Dilp8^{MI00727}$ / +

Bx-Gal4 / +; UAS-Dcr2 / UAS-BrZ2; $Dilp8^{MI00727}$ / +

Bx-Gal4 / +; UAS-Dcr2 / UAS-BrZ3; $Dilp8^{MI00727}$ / +

Bx-Gal4 / +; UAS-Dcr2 / UAS-BrZ4; $Dilp8^{MI00727}$ / +

C: Bx-Gal4 / +; UAS-Eiger / +; $Dilp8^{MI00727}$ / UAS-LacZ.NZ

D: Bx-Gal4 / +; UAS-Eiger / UAS-BrZ1; $Dilp8^{MI00727}$ / +

E: Bx-Gal4 / +; UAS-Eiger / UAS- BrZ2; $Dilp8^{MI00727}$ / +

F: Bx-Gal4 / +; UAS-Eiger / UAS- BrZ3; $Dilp8^{MI00727}$ / +

G: Bx-Gal4 / +; UAS-Eiger / UAS- BrZ4; $Dilp8^{MI00727}$ / +

H: Bx-Gal4 / +; UAS-Eiger / UAS-$Br^{RNAi}$; $Dilp8^{MI00727}$ / +

I-J: genotypes in Fig.S7 C-H

Figure S9

B: AP1-GFP/ +; rn-Gal4, tub-$Gal80^{ts}$ / UAS-LacZ.NZ

C: AP1-GFP / UAS-BrZ1; rn-Gal4, tub-$Gal80^{ts}$ / +

D: AP1-GFP / UAS-BrZ2; rn-Gal4, tub-$Gal80^{ts}$ / +

E: AP1-GFP / UAS-BrZ4; rn-Gal4, tub-$Gal80^{ts}$ / +

Figure S10

B-C: BRV118-GFP / +

D-E: Bx-Gal4 / +; BRV118-GFP / +; UAS-Dcr2 / UAS-LacZ.NZ

Bx-Gal4 / +; BRV118-GFP / UAS-EcR.A$^{W650A}$; UAS-Dcr2 / +

Bx-Gal4 / +; BRV118-GFP / UAS-Br$^{RNAi}$; UAS-Dcr2 / +

Bx-Gal4 / +; BRV118-GFP / UAS-BrZ1; UAS-Dcr2 / +

Bx-Gal4 / +; BRV118-GFP / UAS-BrZ2; UAS-Dcr2 / +

Bx-Gal4 / +; BRV118-GFP / UAS- BrZ4; UAS-Dcr2 / +

Figure S11

A-C: w$^{1118}$ and Dilp8$^{MI00727}$ / Dilp8$^{MI00727}$

D: w$^{1118}$

**Supplementary Figures**

**Figure S1.**

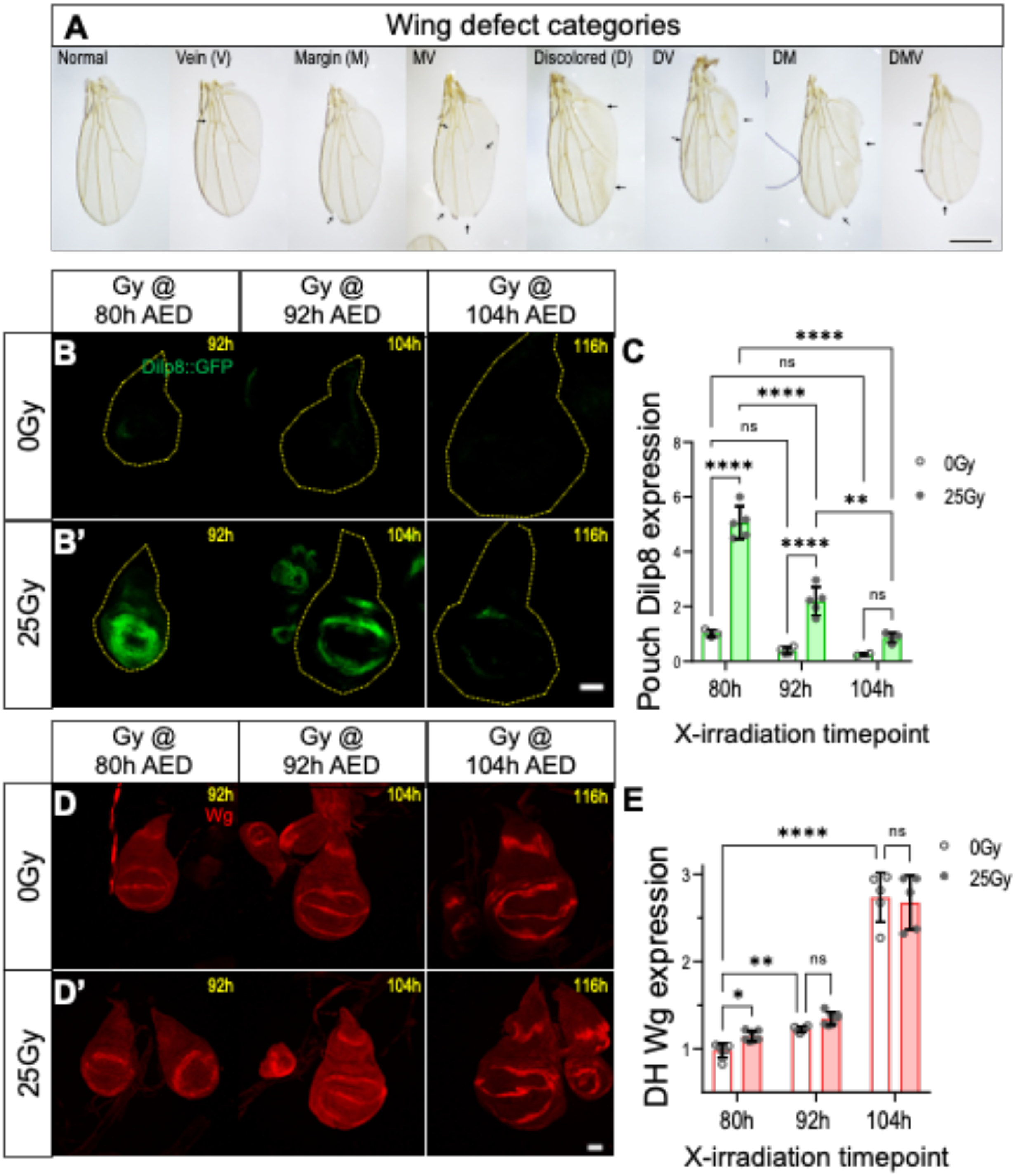

**Figure S1.** Regeneration is restricted as larval development progresses. **(A)** Examples of wings of each damage category that were used in the quantifications in Fig. 1 and Supplemental Fig. 2. Scale=500cm. **(B)** Representative images of wing imaginal discs expressing Dilp8 in undamaged (B) and damaged tissues (B'). Tissues were isolated 12hrs after the damage timepoint (indicated in the figures). Cells expressing Dilp8 are identified by the expression of GFP from the *Dilp8::GFP* transgene construct. Little to no *dilp8* is expressed in the undamaged tissues (B – 80h, 92h, and 104h). In damaged tissues, we see that GFP expression decreases with developmental progression (B' – 80h, 92h, and 104h). Tissues show loss of Dilp8 expression by 104h AED, the Regeneration Restriction Timepoint (RRT). The yellow dotted line indicates tissue area. Scale=50um. **(C)** Quantification of relative *Dilp8::GFP* expression in wing pouch; normalized to GFP expression in undamaged-92h AED tissues. **$p<0.01$, ****$p<0.0001$, two-way ANOVA with Tukey's test. **(D)** Representative images of wingless (Wg) expression in undamaged (D) and damaged tissues (D'). Tissues were isolated 12hrs after the damage timepoint (indicated in the figures). Wg expression increases in the hinge and decreases in the margin following early damage. A similar but less visible increase in hinge Wg is seen following damage at 92h AED. Past the RRT, there is no significant change in Wg expression following damage. Scale=50um. **(E)** Quantification of relative Wg expression in Dorsal Hinge (DH); normalized to DH Wg expression in undamaged-92h AED tissues. *$p<0.05$, **$p<0.01$, ****$p<0.0001$, two-way ANOVA with BK&Y comparisons test

**Figure S2.**

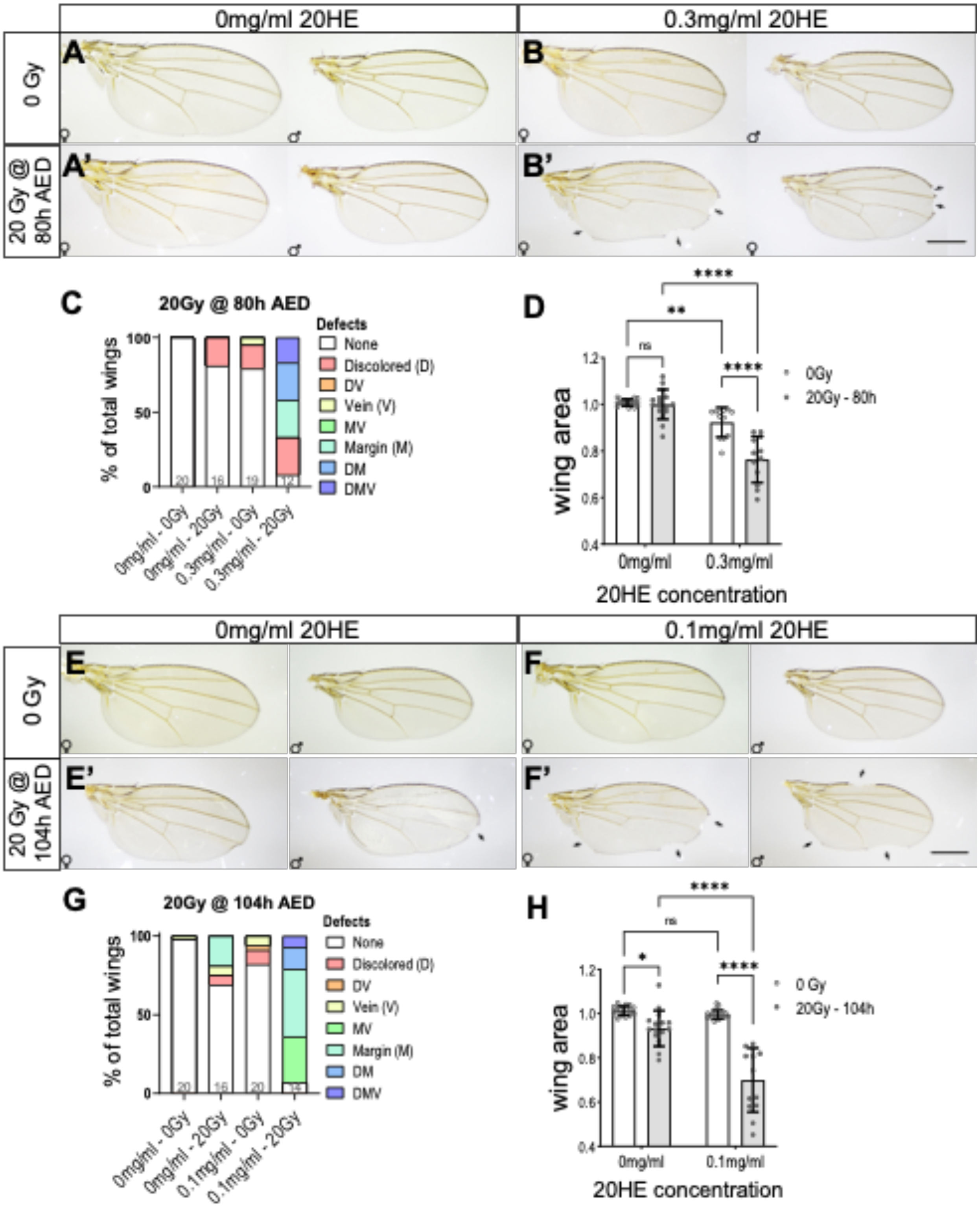
0mg/ml 20HE
0.3mg/ml 20HE
0 Gy
20 Gy @ 80h AED
A
B
A'
B'
C
20Gy @ 80h AED
% of total wings
Defects
None
Discolored (D)
DV
Vein (V)
MV
Margin (M)
DM
DMV
0mg/ml - 0Gy
0mg/ml - 20Gy
0.3mg/ml - 0Gy
0.3mg/ml - 20Gy
D
wing area
0Gy
20Gy - 80h
0mg/ml
0.3mg/ml
20HE concentration
0mg/ml 20HE
0.1mg/ml 20HE
0 Gy
20 Gy @ 104h AED
E
F
E'
F'
G
20Gy @ 104h AED
% of total wings
Defects
None
Discolored (D)
DV
Vein (V)
MV
Margin (M)
DM
DMV
0mg/ml - 0Gy
0mg/ml - 20Gy
0.1mg/ml - 0Gy
0.1mg/ml - 20Gy
H
wing area
0 Gy
20Gy - 104h
0mg/ml
0.1mg/ml
20HE concentration

**Figure S2.** Ecdysone feeding exacerbates damage phenotypes. **(A-B)** Representative images of adult female and male wings of larvae fed ethanol (0mg/ml 20HE) that were either left undamaged (A) or damaged early 20Gy@80h AED (A') and larvae fed 0.3mg/ml 20HE that were either left undamaged (B), or damaged early (B'). Black arrows indicate wing defects. Scale = 500cm. **(C)** Percentage of adult wings that show individual defects or combinations of defects increases following ecdysone feeding. The graph shows the percentage of defective adult wings following no damage (0Gy) and early damage (20Gy-80h) with 0mg/ml or 0.3mg/ml 20HE feeding. **(D)** Quantification of adult wing size following 0Gy and 20Gy-80h with 0mg/ml and 0.3mg/ml 20HE feeding. Data normalized to undamaged wing size of respective sex. **p<0.01, ****p<0.0001, one-way ANOVA with Tukey's test. **(E-F)** Representative images of adult female and male wings of larvae fed ethanol (0mg/ml 20HE) that were either left undamaged (E) or damaged late 20Gy@104h AED (E') and larvae fed 0.1mg/ml 20HE that were either left undamaged (F) or damaged late (F'). Black arrows indicate areas of malformation on late damage wings. Percentage of wings showing malformations indicated on the right. Scale = 500cm. **(G)** Percentage of adult wings that show individual defects or combinations of defects increases following ecdysone feeding. The graph shows the percentage of defective adult wings following no damage (0Gy) and late damage (20Gy-104h) with 0mg/ml or 0.1mg/ml 20HE feeding. **(H)** Quantification of adult wing size following no damaged and early damaged with 0mg/ml or 0.3mg/ml 20HE feeding. Data normalized to undamaged wing size of respective sex. *p<0.05, ****p<0.0001, one-way ANOVA with Tukey's test.

**Figure S3.**

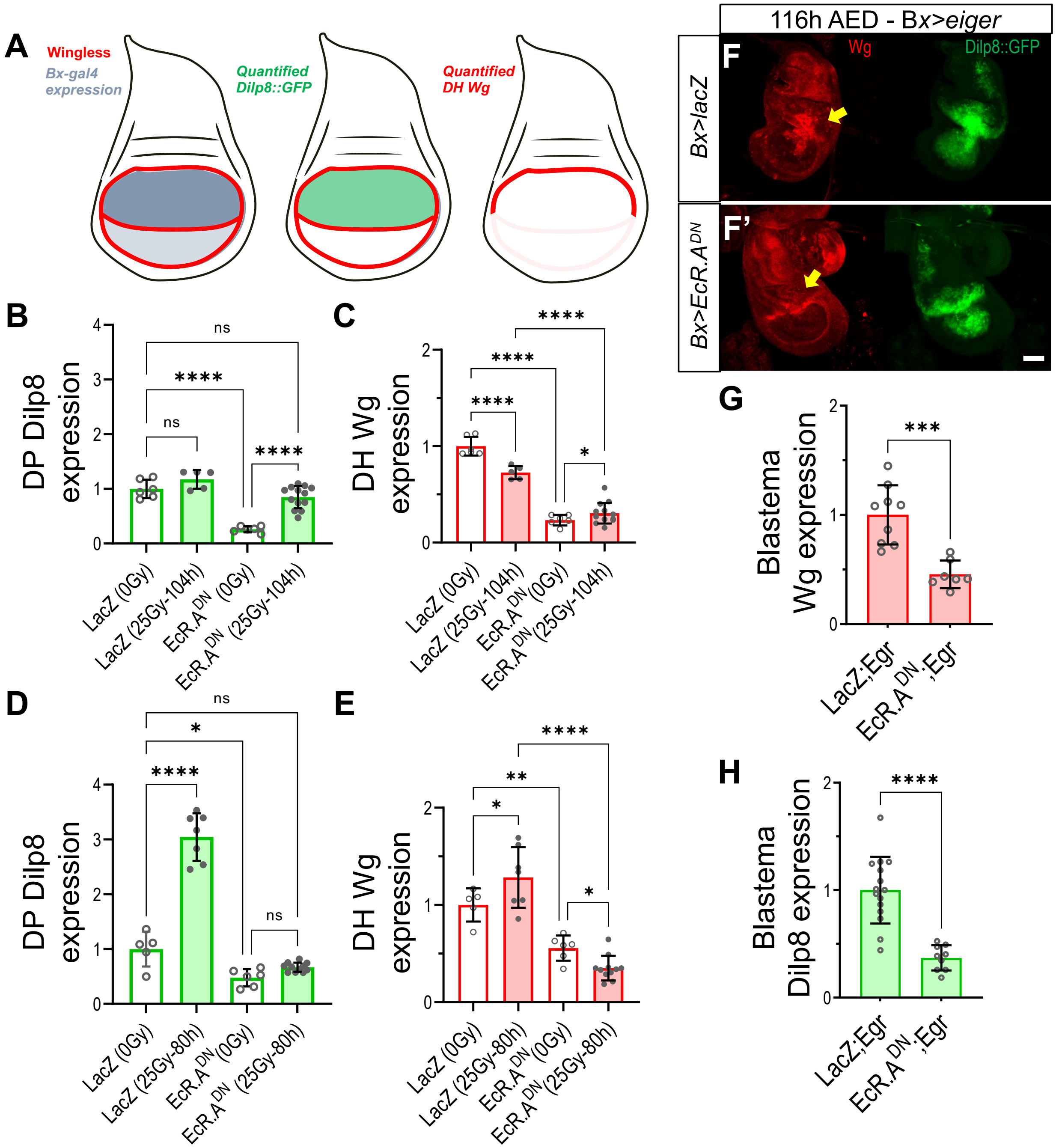
A
Wingless
Bx-gal4 expression
Quantified Dilp8::GFP
Quantified DH Wg
B
DP Dilp8 expression
LacZ (0Gy)
LacZ (25Gy-104h)
EcR.A^DN (0Gy)
EcR.A^DN (25Gy-104h)
C
DH Wg expression
D
DP Dilp8 expression
LacZ (0Gy)
LacZ (25Gy-80h)
EcR.A^DN (0Gy)
EcR.A^DN (25Gy-80h)
E
DH Wg expression
116h AED - Bx>eiger
F
F'
Bx>lacZ
Bx>EcR.A^DN
Wg
Dilp8::GFP
G
Blastema Wg expression
LacZ;Egr
EcR.A^DN;Egr
H
Blastema Dilp8 expression

**Figure S3.** Ecdysone signaling regulates Wg and Dilp8 expression following X-irradiation and *eiger*-induced damage. **(A)** Representative figure of wing disc showing the area of *Bx-gal4* expression in the wing pouch (A). *Bx-gal4* was used in all the wing disc UAS driven overexpression experiments. Quantification of Dilp8 expression changes was done in only the area of highest Bx-gal4 expression (A'). Quantification of Wg was focused on the dorsal hinge, DH (A''). We traced the Wg expression from the anterior margin-hinge intersection to the posterior margin-hinge intersection. **(B-C)** Quantification of dorsal pouch Dilp8::GFP expression (B) and DH Wg expression (C) in 116h AED late damaged (25Gy-104h) and undamaged (0Gy), *Bx>lacZ* and *Bx>EcR.A*$^{DN}$ wing imaginal discs. Quantification normalized to *lacZ (0Gy)*, *p<0.05, ****p<0.0001, one-way ANOVA with Tukey's test (Dilp8) and DYK (Wg) multiple comparisons tests. **(D-E)** Quantification of Dilp8::GFP expression in the dorsal wing pouch (D) and dorsal hinge Wg expression (E) in 92h AED early damaged (25Gy-80h) and undamaged (0Gy), *Bx>lacZ* and *Bx>EcR.A*$^{DN}$ wing imaginal discs. Quantification normalized to *lacZ (0Gy)*, *p<0.05, **p<0.01, ****p<0.0001, one-way ANOVA with Tukey's test (Dilp8) and DYK (Wg) tests. **(F-H)** Representative images of Wg and Dilp8 (Dilp8::GFP) expression in eiger damaged tissues *(Bx>eiger*). The eiger damaged tissues co-expressed *lacZ* (F) and *EcR.ADN* (F'). Yellow arrows indicate the area of eiger expression (the regeneration blastema). Wg and Dilp8 in the blastema are quantified in (G) and (H), respectively. Loss of ecdysone signaling leads to decreased Wg and Dilp8 expression in the blastema. ***p<0.001, ****p<0.0001, Mann Whitney t-test.

**Figure S4.**

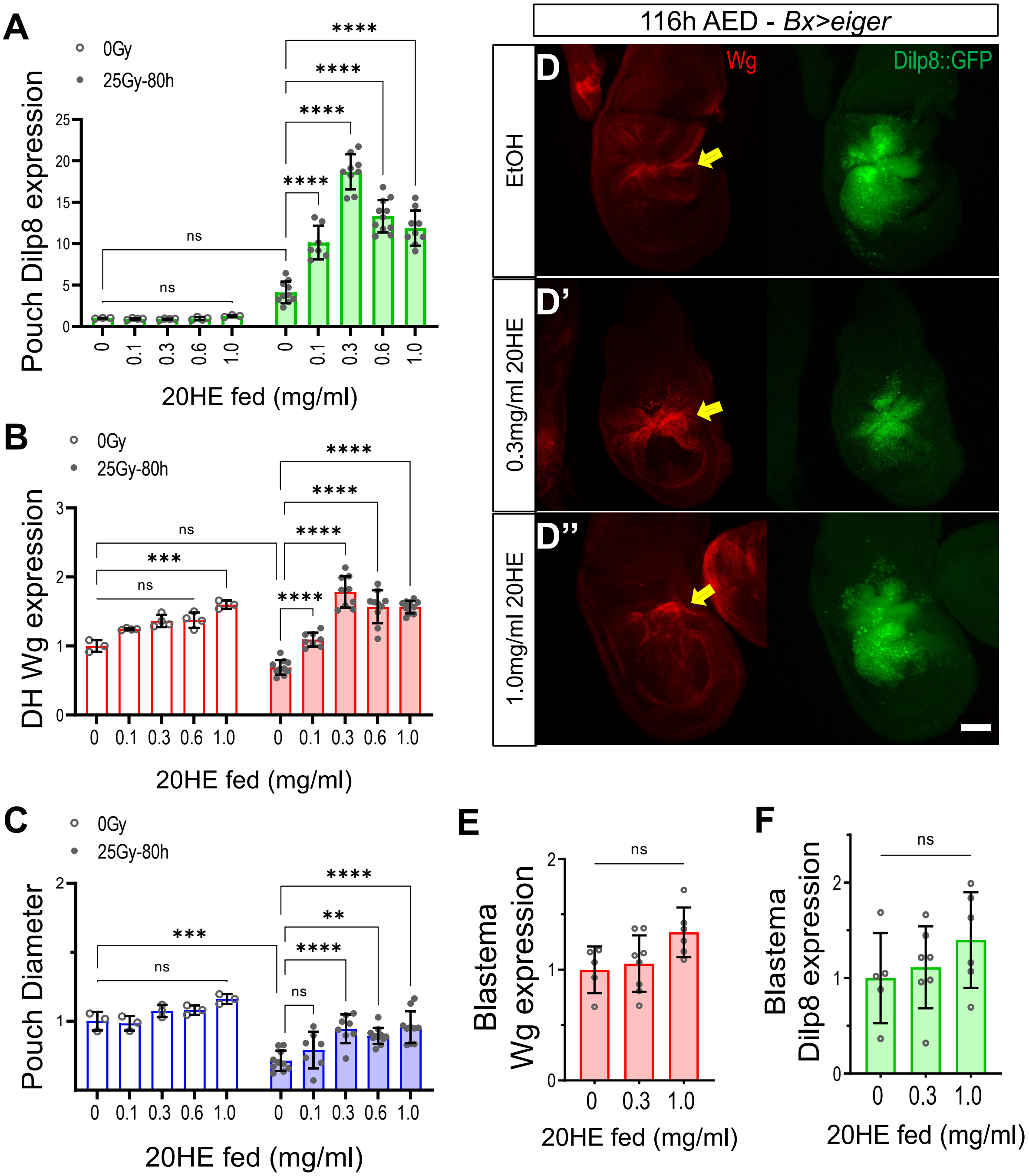
A
0Gy
25Gy-80h
Pouch Dilp8 expression
20HE fed (mg/ml)
ns
****
B
DH Wg expression
***
C
Pouch Diameter
**
116h AED - Bx>eiger
D
D'
D''
Wg
Dilp8::GFP
EtOH
0.3mg/ml 20HE
1.0mg/ml 20HE
E
Blastema Wg expression
F
Blastema Dilp8 expression

**Figure S4.** Increasing systemic ecdysone levels promotes regenerative activity. **(A-C)** Quantification of relative regenerative activity using fold change of Dilp8::GFP expression in the wing pouch (A), dorsal hinge (DH) Wg expression (B), and pouch diameter (C) in 20HE fed and early damaged $w^{1118}$ wing imaginal discs. All quantifications normalized to undamaged tissues 0mg/ml 20HE tissues, **$p<0.01$, ***$p<0.001$, ****$p<0.0001$, two-way ANOVA with Tukey's test. **(D-F)** Representative images of Wg and Dilp8 expression in eiger damaged tissues *(Bx>eiger*) of larvae fed ethanol-0mg/ml (D), 0.3mg/ml (D'), and 1.0mg/ml (D'') of 20HE. Yellow arrows indicate the area of *eiger* expression (regeneration blastema). Wg and Dilp8 in the blastema are quantified in (E) and (F), respectively. Loss of ecdysone signaling leads to decreased Wg and Dilp8 expression in the blastema. Data normalized to 0mg/ml tissues, One-way ANOVA with Tukey's test.

**Figure S5.**

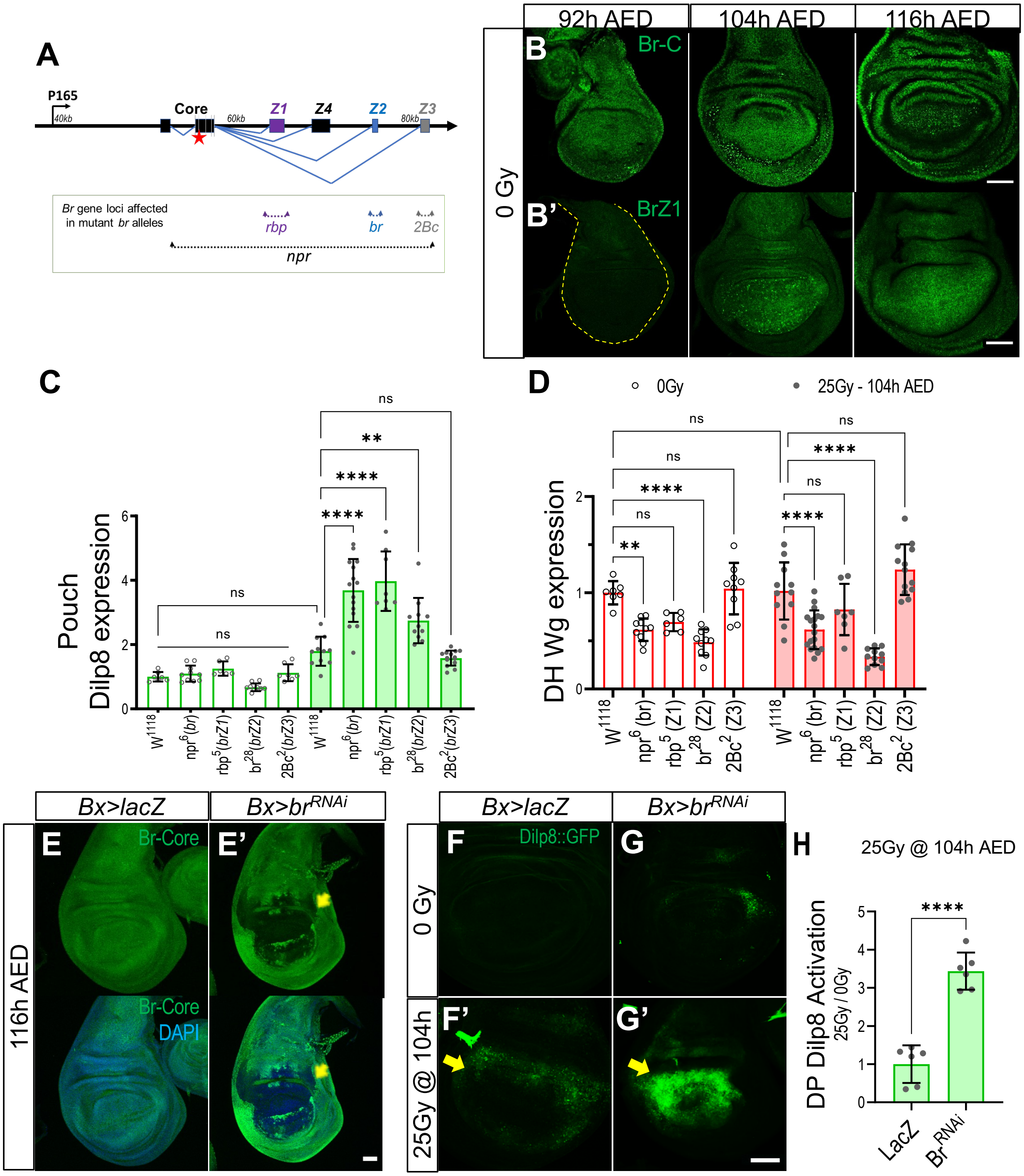

A
P165
40kb
Core
60kb
Z1
Z4
Z2
80kb
Z3
Br gene loci affected in mutant br alleles
rbp
br
2Bc
npr
92h AED
104h AED
116h AED
0 Gy
B
Br-C
B'
BrZ1
C
Pouch Dilp8 expression
D
0Gy
25Gy - 104h AED
DH Wg expression
ns
**
****
Bx>lacZ
Bx>br^RNAi
116h AED
E
E'
Br-Core
DAPI
F
G
F'
G'
Dilp8::GFP
0 Gy
25Gy @ 104h
H
25Gy @ 104h AED
DP Dilp8 Activation
25Gy / 0Gy
LacZ
Br^RNAi

**Figure S5.:** Broad isoforms are expressed in the wing disc and necessary to suppress regenerative activity at the RRT. **(A)** Simplified representation of the *br* gene, starting from P120 promoter (0kb - not included in the diagram) to the end of the *br*-encoding locus. The diagram shows alternate splicing targets for br isoforms and the complementation of their mutant alleles: *npr* (full *br* gene), *rbp (Z1)*, *br (Z2),* and *2Bc (Z3)*. Red star indicates UAS-$br^{RNAi}$ (FBgn0283451) target. **(B)** Representative time-course images of Br (A – Br-core (Br-C)) and BrZ1 (A’) expression in undamaged tissue. IF staining validates BrZ1 expression in wing imaginal discs seen in the Western blot (Fig 4a: 70-75kD band). **(C-D)** Quantification of regenerative activity in 116h AED tissues using pouch Dilp8::GFP expression in the (C) and dorsal hinge (DH) Wg expression (D) in $w^{1118}$ and *br* mutants wing imaginal discs. All quantifications normalized to undamaged $w^{1118}$, **p<0.01, ****p<0.0001, two-way ANOVA with Tukey’s test. **(E-E’)** Representative images of *br* knockdown using *Bx>*$br^{RNAi}$ (E’) vs. control *Bx>lacZ* (E). Tissues stained with br-core antibody and DAPI. Yellow arrows indicate the area of *br* knockdown. Scale bar=50um. **(F-H)** Representative images of pouch Dilp8::GFP 116h AED undamaged (F-G), and late damaged - 25Gy @ 104h AED (F’-G’) wing imaginal discs. Primary area of expression indicated by yellow arrows. Scale bar=50um Tissues are expressing *Bx>lacZ* as a control (F-F’) or *Bx>*$br^{RNAi}$ (G-G’), Dilp8::GFP expression fold change (H) is calculated by normalizing to their respective 116h AED undamaged controls, ****p<0.0001, Mann-Whitney t-test.

**Figure S6.**

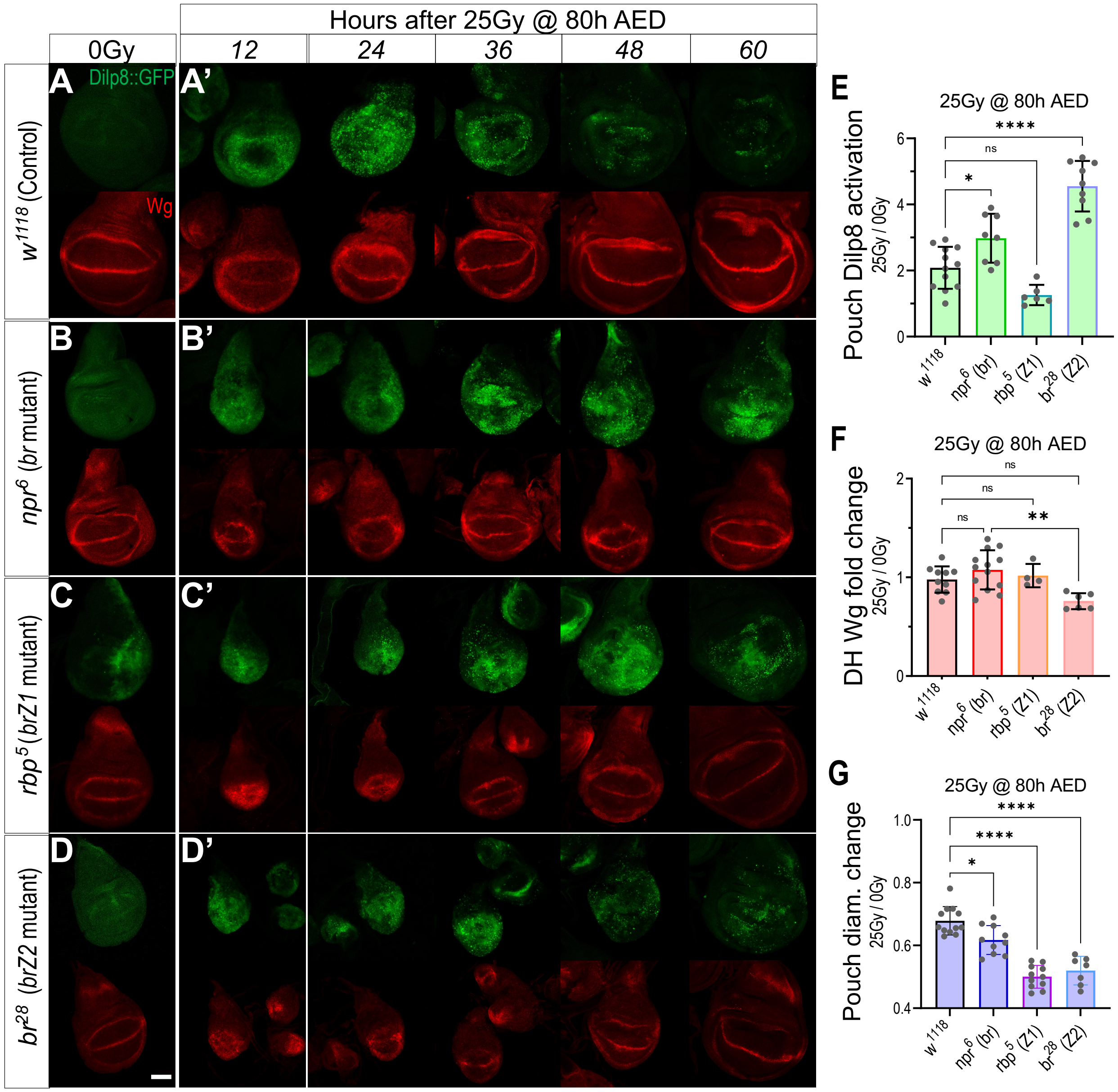
Hours after 25Gy @ 80h AED
0Gy
12
24
36
48
60
A
Dilp8::GFP
A'
Wg
w1118 (Control)
B
B'
npr6 (br mutant)
C
C'
rbp5 (brZ1 mutant)
D
D'
br28 (brZ2 mutant)
E
25Gy @ 80h AED
Pouch Dilp8 activation
25Gy / 0Gy
****
ns
*
w1118
npr6 (br)
rbp5 (Z1)
br28 (Z2)
F
25Gy @ 80h AED
DH Wg fold change
25Gy / 0Gy
ns
ns
ns
**
G
25Gy @ 80h AED
Pouch diam. change
25Gy / 0Gy
****
****
*

**Figure S6:** Broad isoforms regulate the timing and duration of the regenerative response – Part A. **(A-D)** Time-course images of Dilp8::GFP (green) and Wg (red) expression following early damage (25Gy @ 80h AED). Representative undamaged tissues, 12hr after damage timepoint, are shown in A-D. Damaged tissue isolated in 12hr intervals are shown in A'-D'. Tissues used are control – $w^{1118}$ (A-A'), and *br* mutants: full *br* mutant - $npr^6$ (B-B'), *brZ1* mutant - $rbp^5$ (C-C'), and *brZ2* mutant - $br^{28}$ (D-D'). Scale bar=50um. **(E-G)** Fold change in Dilp8 expression (E), Wg expression (F), and pouch diameter (G) following early damage in $w^{1118}$ controls and *br* mutants: **p<0.01, ****p<0.0001, one-way ANOVA with Tukey's test.

**Figure S7.**

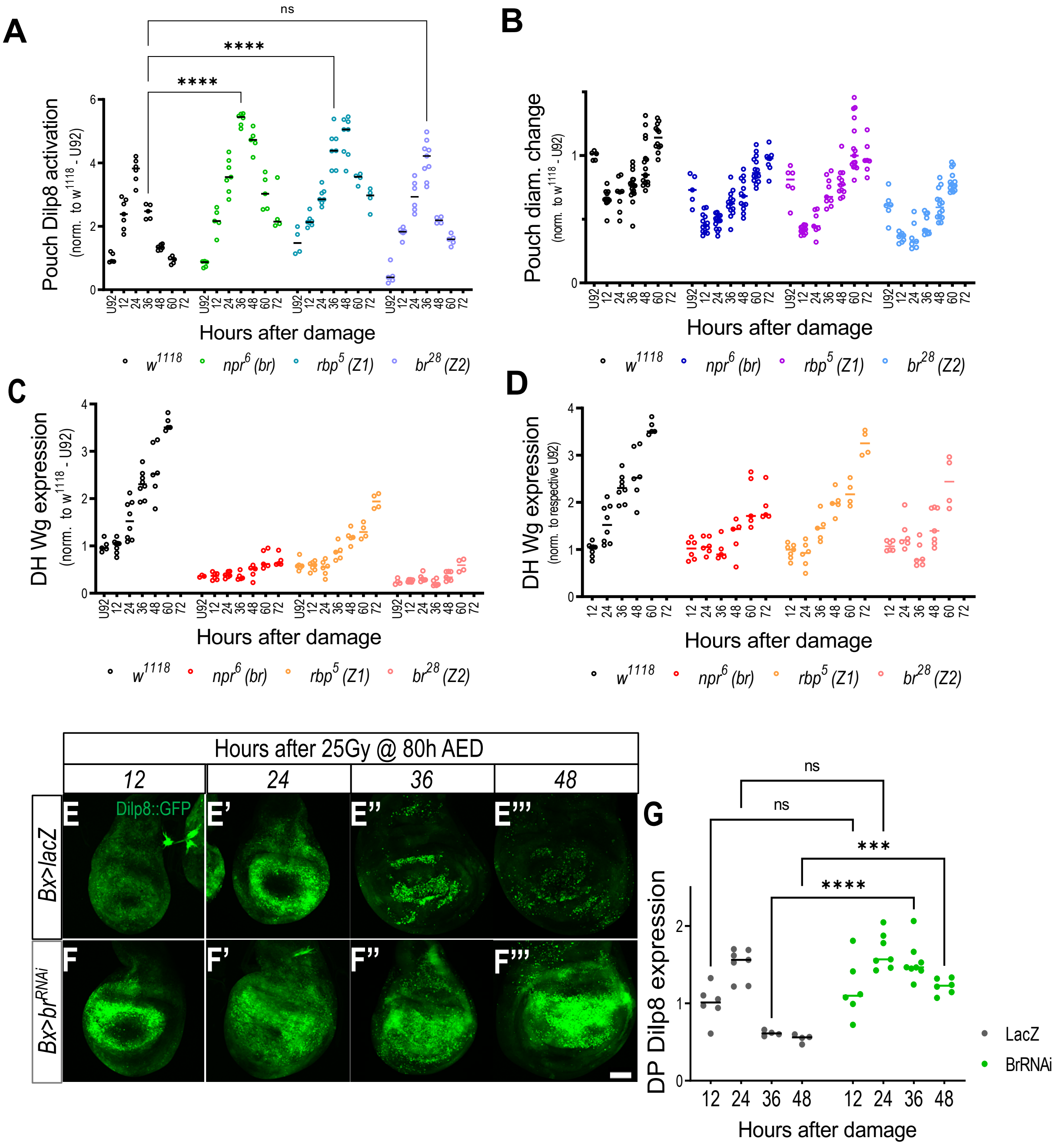
A
B
C
D
Pouch Dilp8 activation
(norm. to w1118 - U92)
Pouch diam. change
DH Wg expression
(norm. to respective U92)
Hours after damage
ns
****
w1118
npr6 (br)
rbp5 (Z1)
br28 (Z2)
Hours after 25Gy @ 80h AED
12
24
36
48
Bx>lacZ
Bx>brRNAi
Dilp8::GFP
E
E'
E''
E'''
F
F'
F''
F'''
G
DP Dilp8 expression
***
LacZ
BrRNAi

**Figure S7.** Broad isoforms regulate the timing and duration of the regenerative response – Part B. **(A-D)** Relative expression of Dilp8::GFP (A), pouch diameter (B), and dorsal hinge (DH) Wg expression (C-D). U92's (undamaged 92h AED) are also included in the graph. All data normalized to $w^{1118}$ U92. 72h *npr*$^{6}$ and *rbp*$^{5}$ tissues are also included in the graph. **(E-F)** Representative time-course images of Dilp8 expression following early damage (25Gy @ 80h AED) in *Bx>lacZ* (E-E''') and *Bx>br*$^{RNAi}$ (F-F''') tissues. Tissues were isolated in 12-hour intervals following damage. **(G)** Quantification of Dilp8::GFP expression in the dorsal pouch (DP), normalized to expression in *Bx>lacZ* tissues isolated 12 hours after damage. ***p<0.001, ****p<0.0001, two-way ANOVA with Tukey's test.

Figure S8.

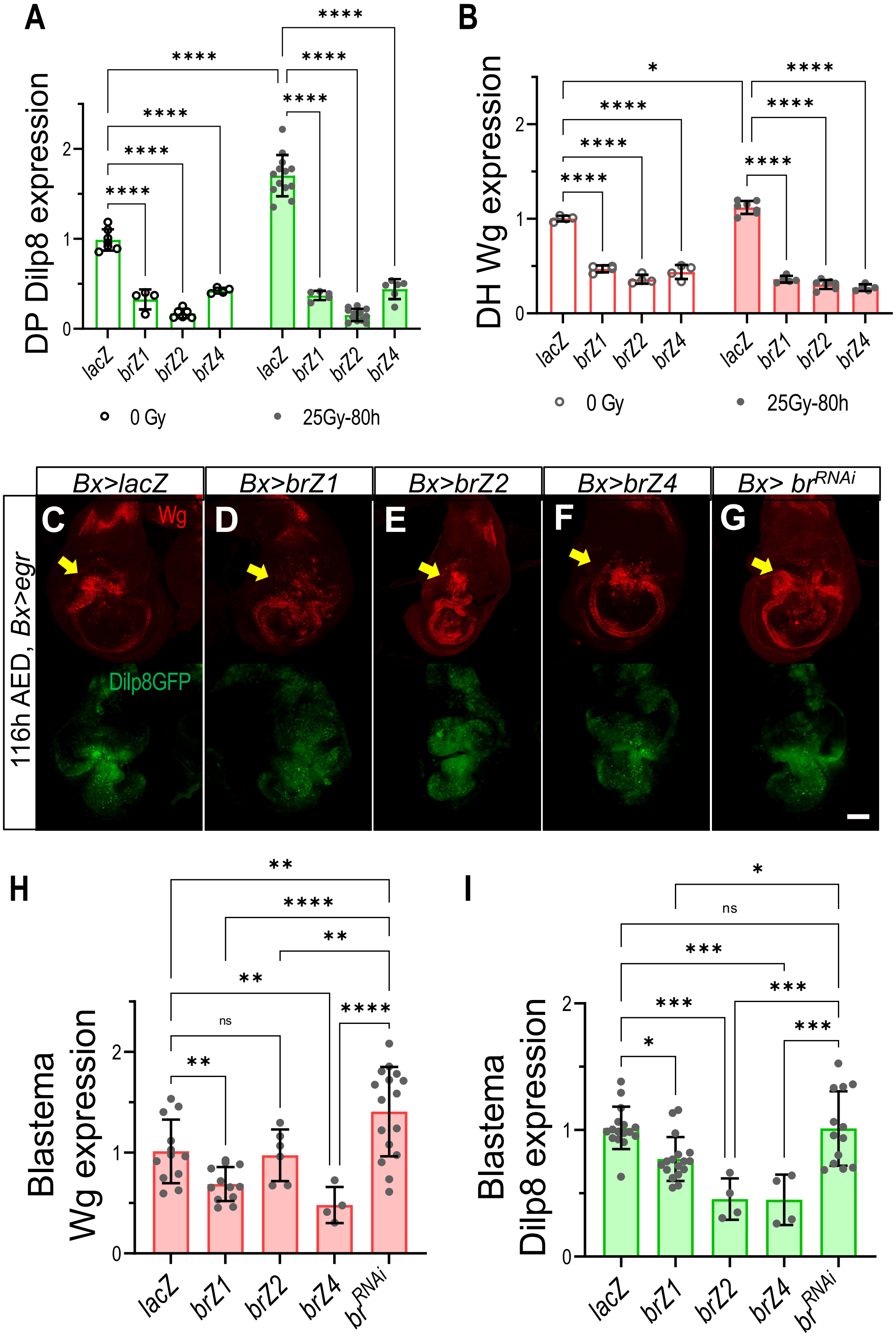

A
DP Dilp8 expression
lacZ
brZ1
brZ2
brZ4
0 Gy
25Gy-80h
B
DH Wg expression
C
D
E
F
G
Bx>lacZ
Bx>brZ1
Bx>brZ2
Bx>brZ4
Bx> br^RNAi
116h AED, Bx>egr
Wg
Dilp8GFP
H
Blastema Wg expression
I
Blastema Dilp8 expression
ns

**Figure S8.** Broad isoforms are sufficient to suppress regenerative activity following X-irradiation and *eiger*-induced damage. **(A-B)** Quantification of regenerative activity in 92h AED tissues using dorsal pouch Dilp8::GFP expression in the (A) and dorsal hinge (DH) Wg expression (B) in undamaged and early damaged wing discs and *br* mutants wing imaginal discs. All quantifications normalized to undamaged $w^{1118}$, ****p<0.0001, **p<0.01, two-way ANOVA with Tukey's test. **(C-G)** Representative images of Wg and Dilp8 (Dilp8::GFP) expression in eiger damaged tissues *(Bx>eiger)*. The eiger damaged tissues co-expressed *lacZ* (C), *brZ1* (D), *brZ2* (E), *brZ4* (F), and $br^{RNAi}$ (G). **(H, I)** Broad isoforms suppress Wg expression in the pouch. Wg and Dilp8 expression in the blastema. Broad knockdown leads to increased Wg expression in the blastema. Yellow arrows indicate the area of *eiger* expression (the regeneration blastema). Expression of Wg and Dilp8 in the blastema is quantified in (H) and (I), respectively; normalized to *lacZ* controls. *p<0.05, **p<0.01, ***p<0.001, ***p<0.001, ****p<0.0001, one-way ANOVA with Tukey's test.

**Figure S9.**

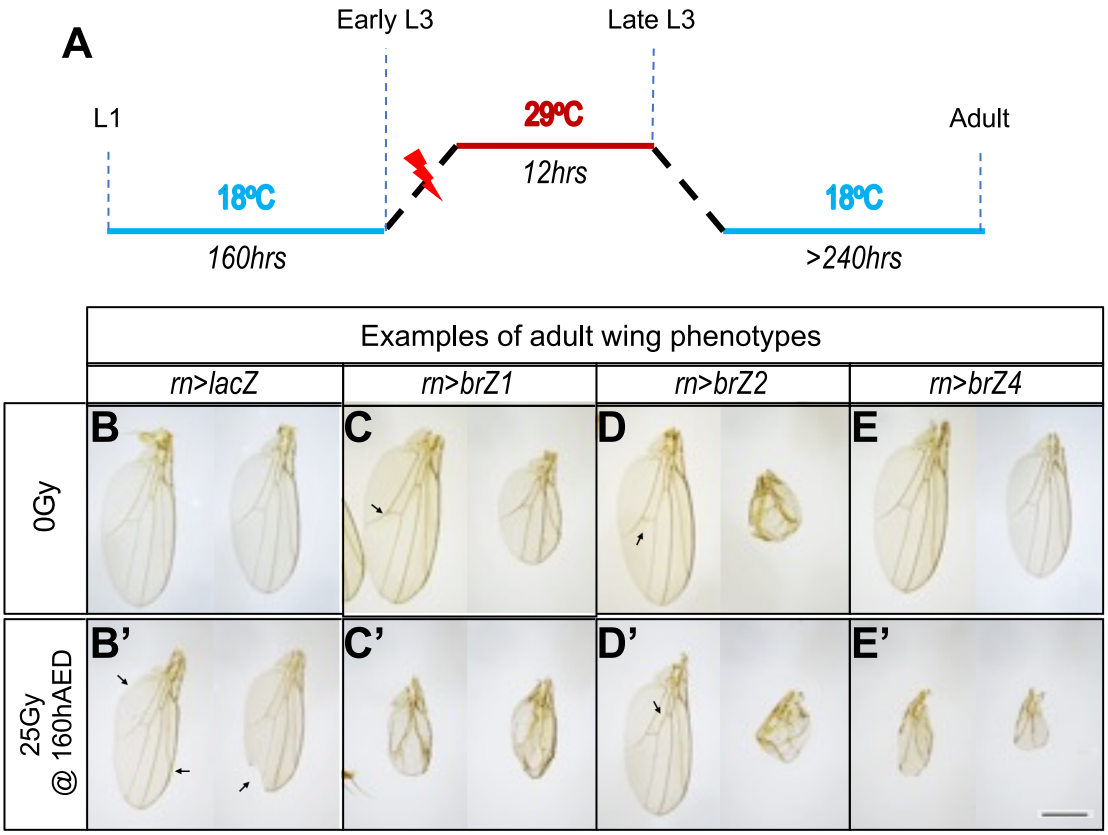

A
Early L3
Late L3
L1
29ºC
Adult
12hrs
18ºC
18ºC
160hrs
>240hrs
Examples of adult wing phenotypes
rn>lacZ
rn>brZ1
rn>brZ2
rn>brZ4
B
C
D
E
0Gy
B'
C'
D'
E'
25Gy
@ 160hAED

**Figure S9.** Broad isoforms limit regeneration. **(A)** Schematic of transient Br isoform overexpression experiment. rn-gal4 driven expression of UAS-constructs was limited to L3 using temperature-sensitive tubulin-gal80 (tub-gal80$^{ts}$). Stocks were maintained at 18$^{o}$C until early L3, then irradiated (red lightning – 25Gy) or left unirradiated (0Gy). Following a 12hr incubation period at 29$^{o}$C, the larvae were returned to 18$^{o}$C until the adult stage. **(B-F)** Examples of adult wings arising from 0Gy (B-F) and 25Gy (B'-F') damaged larvae following overexpression of control – LacZ (B-B') and Br isoforms: BrZ1 (C-C'), BrZ2 (D-D'), BrZ3 (E-E'), and BrZ4 (F-F'). Black arrows indicate defects in wings that otherwise appear normal. Examples of normal wings can be found in (B). Scale=500cm **(G)** Percentage of adult wings that show individual defects or combinations of defects increases following ecdysone feeding. The graph shows the percentage of defective adult wings following no damage (0Gy) and late damage (25Gy-104h) with 0mg/ml or 0.1mg/ml 20HE feeding. **(H)** Quantification of adult wing size following 0Gy and 25Gy after transient expression of broad isoforms. Data normalized to undamaged lacZ wings of respective sex. One-way ANOVA with Tukey's test, *$p<0.05$, ***$p<0.001$, ****$p<0.0001$.

**Figure S10.**

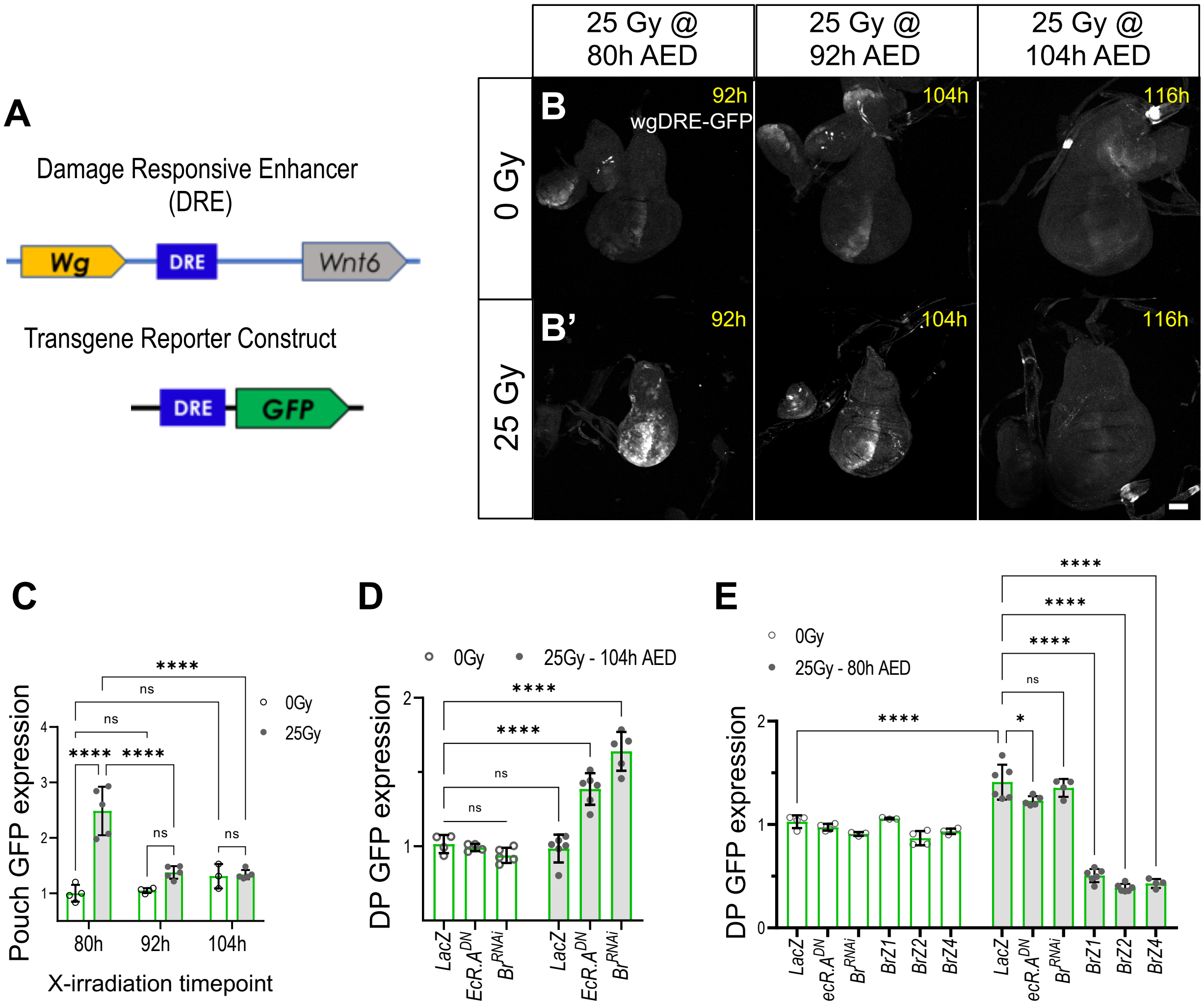
A
Damage Responsive Enhancer
(DRE)
Wg
DRE
Wnt6
Transgene Reporter Construct
DRE
GFP
25 Gy @
80h AED
25 Gy @
92h AED
25 Gy @
104h AED
B
B'
0 Gy
25 Gy
92h
104h
116h
wgDRE-GFP
C
Pouch GFP expression
0Gy
25Gy
80h
92h
104h
X-irradiation timepoint
ns
****
D
0Gy
25Gy - 104h AED
DP GFP expression
LacZ
EcR.A^DN
Br^RNAi
E
0Gy
25Gy - 80h AED
DP GFP expression
LacZ
ecR.A^DN
Br^RNAi
BrZ1
BrZ2
BrZ4
*

**Figure S10.** wgDRE is suppressed at the end of larval development by broad isoforms. **(A)** Simplified representation of the *wg* and *wnt*$^{6}$ gene locus. The diagram shows the location of the *wg*-Damage Responsive Enhancer (DRE aka BRV118) and the *wg*DRE-GFP transgene construct used in B-E below. **(B-C)** Representative images of wgDRE activation in undamaged (B) and damaged tissues (B'). *wg*DRE is slightly activated in undamaged tissue during development (B – 80h, 92h, and 104h). In damaged tissues, *wg*DRE activation is much higher and shows a loss of this damage-induced activation at the RRT (B' – 80h, 92h, and 104h). Quantification of GFP expression (C) is normalized to GFP expression in undamaged 80h AED tissues. ANOVA Brown-Forsythe test with Tukey's test, ****p<0.0001 **(D)** Relative dorsal pouch (DP) *wg*DRE-GFP expression following late damaged control - *Bx>lacZ, Bx>EcR.A*$^{DN}$ and *Bx>br*$^{RNAi}$ overexpressing wing imaginal discs. Quantification normalized to 116h AED undamaged *lacZ* tissues. ****p<0.0001, two-way ANOVA with Tukey's test. **(E)** Relative dorsal pouch (DP) *wg*DRE-GFP expression following early damaged control - *Bx>lacZ* and *br* isoform overexpressing wing imaginal discs. Quantification normalized to 92h AED undamaged *lacZ* tissues. ****p<0.0001, **p<0.01, *p<0.05, two-way ANOVA with Tukey's test.

**Figure S11.**

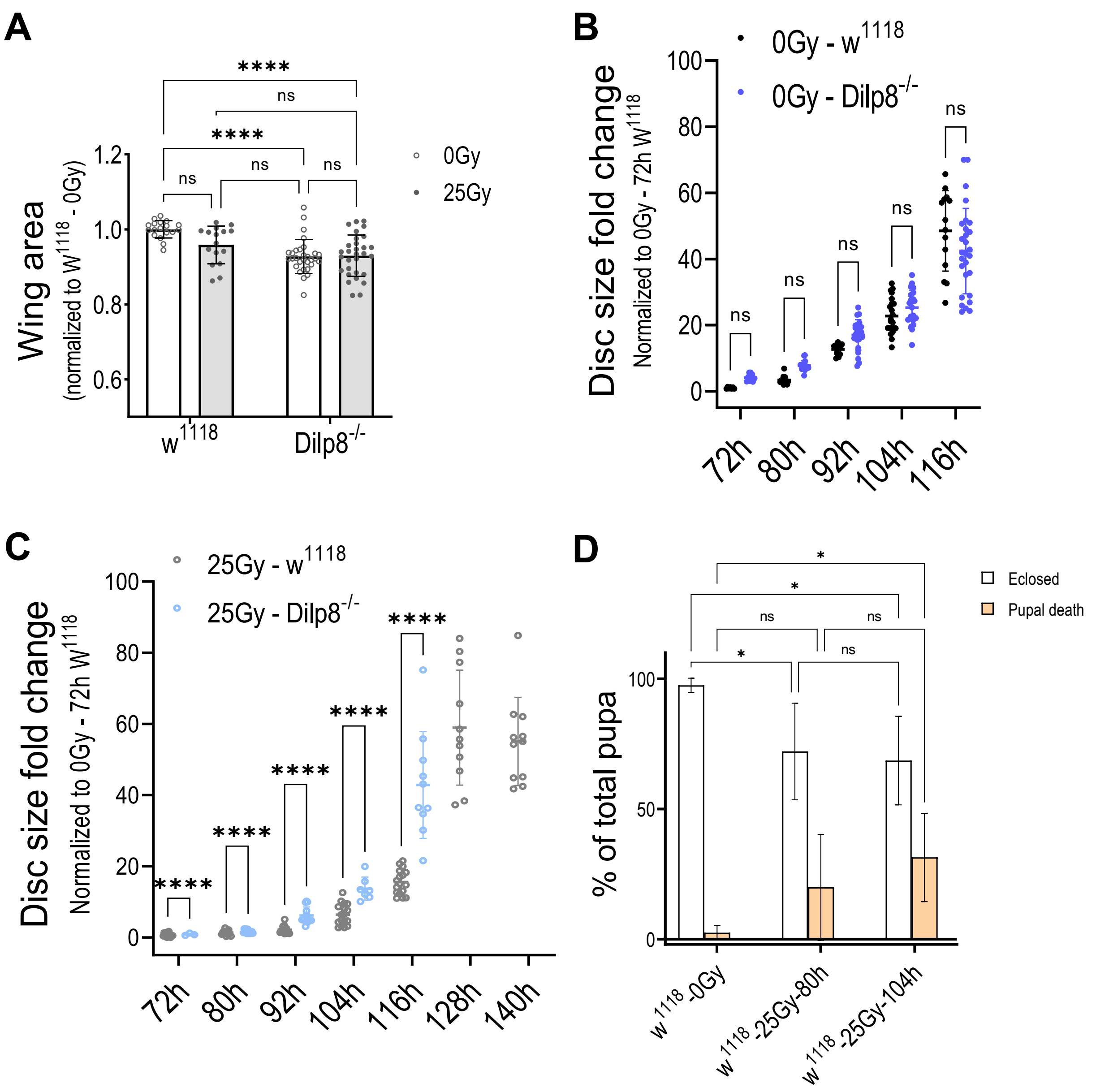

A
Wing area
(normalized to W1118 - 0Gy)
1.2
1.0
0.8
0.6
****
ns
****
ns
ns
ns
0Gy
25Gy
w1118
Dilp8-/-
B
Disc size fold change
Normalized to 0Gy - 72h W1118
0Gy - w1118
0Gy - Dilp8-/-
100
80
60
40
20
0
ns
72h
80h
92h
104h
116h
C
Disc size fold change
Normalized to 0Gy - 72h W1118
25Gy - w1118
25Gy - Dilp8-/-
****
72h
80h
92h
104h
116h
128h
140h
D
% of total pupa
100
50
0
Eclosed
Pupal death
*
ns
w1118 -0Gy
w1118 -25Gy-80h
w1118 -25Gy-104h

**Figure S11. Regeneration dynamics in *dilp8*$^+$ and *dilp8*$^-$ larvae**

**(A)** Quantification of adult wing size for tissue in adult wings following no damage (0Gy) and early damage (25Gy-80h) in $w^{1118}$ and $dilp8^{-/-}$ adults. Size of wing measured in the unit area and normalized to undamaged $w^{1118}$ wing size of respective genotype and sex. 2-way ANOVA with Tukey's test, ****$p<0.0001$. **(B-C)** Quantification of wing disc growth in $w^{1118}$ and $dilp8^{-/-}$ tissues following no damage - 0Gy, (B) and damage - 25Gy-48h AED (C). Data sets normalized to 72h $w^{1118}$ tissues. ***$p<0.001$, ****$p<0.0001$, 2way-ANOVA with Tukey's test. **(D)** Quantification of population viability in $w^{1118}$ following no damage - 0Gy, early damage 25Gy-80h, and late damage 25Gy-104h. *$p<0.05$, two-way ANOVA with Tukey's test.
